# ARID3A coordinates the proliferation-differentiation switch of transit-amplifying cells in the intestine

**DOI:** 10.1101/2023.09.25.559311

**Authors:** Nikolaos Angelis, Anna Baulies, Anna Kucharska, Gavin Kelly, Miriam L Sopena, Stefan Boeing, Vivian S.W. Li

## Abstract

Intestinal stem cells (ISCs) at the crypt base divide and give rise to progenitor cells that have the capacity to proliferate and differentiate into various mature epithelial cell types in the transit-amplifying (TA) zone. Here, we identified the transcription factor ARID3A as a novel regulator of intestinal epithelial cell proliferation and differentiation at the TA compartment. We show that ARID3A forms an expression gradient from villus tip to the early progenitors at the crypts mediated by TGF-β and WNT signalling. Intestinal epithelial-specific deletion of *Arid3a* reduces proliferation of TA cells. Bulk and single cell transcriptomic analysis shows increased enterocyte differentiation and reduced secretory cells in the *Arid3a* cKO intestine. Interestingly, upper-villus gene signatures of both enterocytes and secretory cells are enriched in the mutant intestine. We find that the enhanced enterocyte differentiation in the *Arid3a* cKO intestine is caused by increased binding of HNF1 and HNF4. Finally, we show that loss of *Arid3a* impairs irradiation-induced regenerative process by altering the dynamics of proliferation and apoptosis. Our findings imply that ARID3A may play a gatekeeping role in the TA compartment to maintain the “just-right” proliferation-to-differentiation ratio for tissue homeostasis and plasticity.

## Introduction

The intestinal epithelium is one of the fastest renewing and regenerating tissues and its high turnover in cell composition is facilitated by Lgr5+ intestinal stem cells (ISCs) residing at the intestinal crypts. ISCs divide to generate a daughter cell that will either self-renew to generate another stem cell or will enter the transit-amplifying (TA) zone for subsequent lineage specification (1–3). Intestinal lineage decision takes place at cell positions +4/+5, where progenitor cells are located (4). These progenitors can be broadly divided into two main subtypes, absorptive and secretory, which are highly plastic and are able to re-acquire stemness for tissue regeneration upon injury (5–7). The stem cell-to-daughter cell transition in the intestinal epithelium is a highly dynamic and plastic process. Maintenance and regulation of the stem cell pool is controlled both by an epithelial cellular niche as well as by the mucosal stromal microenvironment (8).

A variety of signalling cascades has been well-described in regulating ISC maintenance, fate decision and terminal maturation. WNT and NOTCH signalling are the key drivers to maintain stem cell identity at the bottom of the crypt. At the +4/+5 progenitors, NOTCH dictates the initial absorptive versus secretory fate decision: “NOTCH-ON” promotes enterocyte differentiation, whilst “NOTCH-OFF” de-represses the master regulator ATOH1 to drive secretory lineage specification. We have recently identified that the transcription co-repressors MTG8 and MTG16 are also key regulators expressed at the progenitors to facilitate this binary fate decision process by repressing ATOH1 transcription (9). After the initial lineage commitment process at +4/5 cells, these progenitors will continue to proliferate and differentiate at the TA cells in the upper crypts and will eventually exit the cell cycle for terminal differentiation in the villi. Emerging evidence reveals that enterocytes, goblet cells and tuft cells exhibit a broad zonation of their gene expression programme along the crypt-villus axis to facilitate different functions, highlighting the complexity of the cellular differentiation process (10–12). Whilst the initial lineage specification at +4/5 cells has been extensively studied, the molecular control of proliferation and differentiation states at the TA cells has been largely overlooked in the past. Characterising the TA cell regulation will be important for the understanding of intestinal epithelial cell type composition under homeostasis and injury-induced regeneration.

Here, we report the transcription factor ARID3A as a novel regulator of intestinal homeostasis that plays a central role in controlling proliferation and differentiation dynamics of the TA cells. ARID3A forms an expression gradient from villus tip to crypt progenitor cells driven by TGF-b and WNT signalling. Loss of *Arid3a* inhibits TA cell proliferation, perturbs the absorptive versus secretory cell differentiation and increases the expression of villus-tip gene signatures, leading to reduced regenerative capacity upon irradiation-induced injury. Our findings reveal the hitherto unrecognised role of ARID3A in coordinating TA cell proliferation and differentiation in the intestine.

## Results

### *Arid3a* forms an expression gradient from villus tip to crypt progenitor cells

We have previously identified a set of genes that are enriched at the +4/+5 progenitor cells compared to the *Lgr5*+ ISCs (9). Screening for transcription factors that are enriched at the +4/+5 cells identified the A+T Rich Interaction Domain 3a (*Arid3a*) as a putative modulator of intestinal epithelial homeostasis (Figure 1A). Quantitative reverse transcription (qRT-PCR) analysis of Lgr5-GFP sorted cells isolated from a *Lgr5-EGFP-ires-CreERT2* intestinal crypts confirmed the enrichment at the GFP-low progenitors (Figures 1B and S1A). To further probe the localisation of *Arid3a* at the crypt, we performed double RNAscope co-staining of *Arid3a* with *Lgr5* (ISC marker) and *Atoh1* (secretory progenitor and Paneth cell marker). While *Arid3a* showed both overlapping and exclusive staining with *Lgr5* and *Atoh1,* the co-localisation was minimal (Figure 1C). Quantification of the RNAscope data revealed that *Arid3a* not only was enriched at the +4/+5 cells as compared to the ISCs, but its expression was maintained throughout the upper crypt (Figure 1D). Interestingly, expression analysis of ARID3A at both mRNA and protein levels showed a strong expression gradient from the villus tip to early progenitor cells at the crypt (Figures 1E and 1F). To confirm the enrichment of *Arid3a* at the villus compartment, we performed crypt-villus fractionation of mouse proximal small intestinal tissue followed by qRT-PCR analysis of the two compartments. As expected, the stem cell-specific marker *Olfm4* was enriched at the crypt fraction, while the enterocyte marker Alkaline phosphatase (*Alpi*) was enriched at the villus. In accordance with our RNAscope data, *Arid3a* showed a 20-fold upregulation in the villus compared to the crypt (Figure S1B). Stromal expression of *Arid3a* was also detected (Figures 1E and 1F) since *Arid3a* is also expressed in B cells (13, 14).

**Figure 1.**
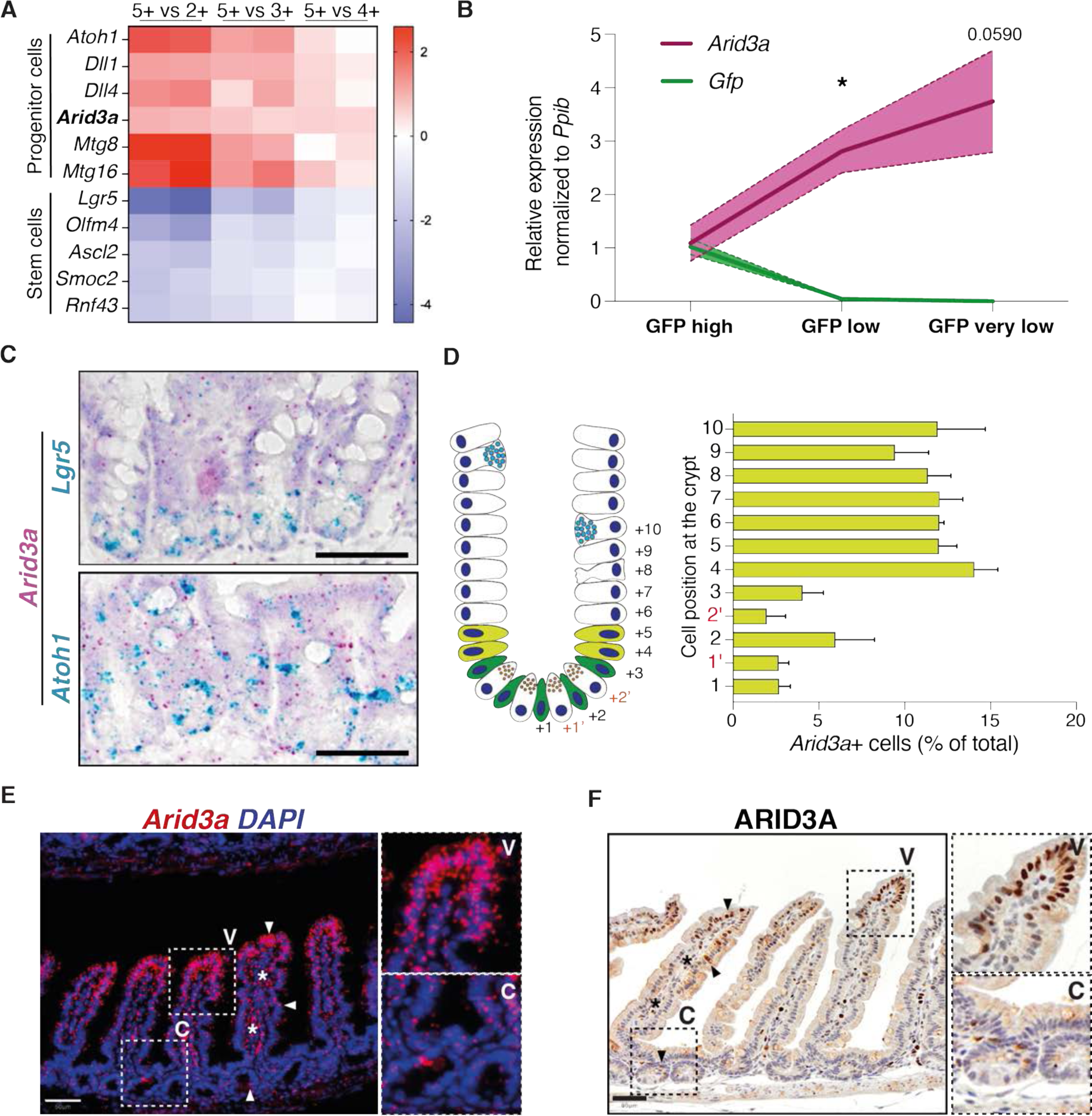
ARID3A follows a gradient of expression across the crypt-villus axis. (A) Re-analysis of differential gene expression between Lgr5-positive stem cells and their immediate progeny (GSE36497) shows enrichment of *Arid3a* at the progenitor cells. (B) qRT-PCR analysis of three FACS sorted populations representing ISCs (GFP high) and progenitor cells (GFP low and GFP very low). Three biologically independent animals (N=3). Data represent mean ± s.e.m. (light colour area) *P<0.05, **P<0.01, ***P<0.001, two-sided t-test. (C) Dual RNAscope of *Arid3a* with *Lgr5* or Atoh1 at WT mouse small intestine. Each staining was performed at three different animals (N=3). (D) Quantification of *Arid3a*+ cells per crypt position. Analysis was performed using the Arid3a/Lgr5 co-stain. (E) RNAscope of *Arid3a* at WT mouse small intestine. Staining was performed at three different animals (N=3). (F) Protein staining of ARID3A at WT mouse small intestine. Staining was performed at three different animals (N=3).

Enterocytes are the most prominent type of intestinal epithelial cells exhibiting characteristic microvilli structures (15). Careful examination of ARID3A protein staining confirmed the expression of ARID3A in brush border-bearing enterocytes (Figure 1F). We further tested if *Arid3a* is also expressed at secretory cells. By combing RNAscope staining of *Arid3a* and immunofluorescent staining to detect the protein levels of Mucin 2 (MUC2), Chromogranin A (CHGA) and Lysozyme (LYZ), we confirmed that *Arid3a* is also expressed in goblet, enteroendocrine and Paneth cells, respectively (Figure S1C).

### WNT and TGF-β regulate expression of *Arid3a* in epithelial cells

Next, we sought to characterise the driver of *Arid3a* expression gradient. *Arid3a* forms an expression gradient from the upper crypt in a pattern opposite to the WNT gradient, implying that WNT signalling may regulate its expression. To test the regulatory role of WNT, *ex vivo* wild-type (WT) mouse intestinal organoids were treated with two different WNT inhibitors, LF3 and LGK974, for 48h. Successful WNT inhibition was confirmed by downregulation of known WNT target genes, such as *Axin2*, *Sox9* and *Cyclin D1* (Figure 2A and S2A). In contrast, *Arid3a* expression was upregulated upon treatment with either of the two inhibitors, suggesting an repressive role of WNT in *Arid3a* expression (Figures 2A and S2A). To validate the negative role of WNT in *Arid3a* expression, we further examined an independent WNT overactivation model using our previously published WNT-high mouse organoids carrying a truncated APC (APC5) (16). Consistent with the observations earlier, *Arid3a* expression was downregulated, while the WNT targets *Axin2* and *Cyclin D1* were significantly increased in the WNT-high Apc5 organoids compared to WT control (Figure 2B).

**Figure 2.**
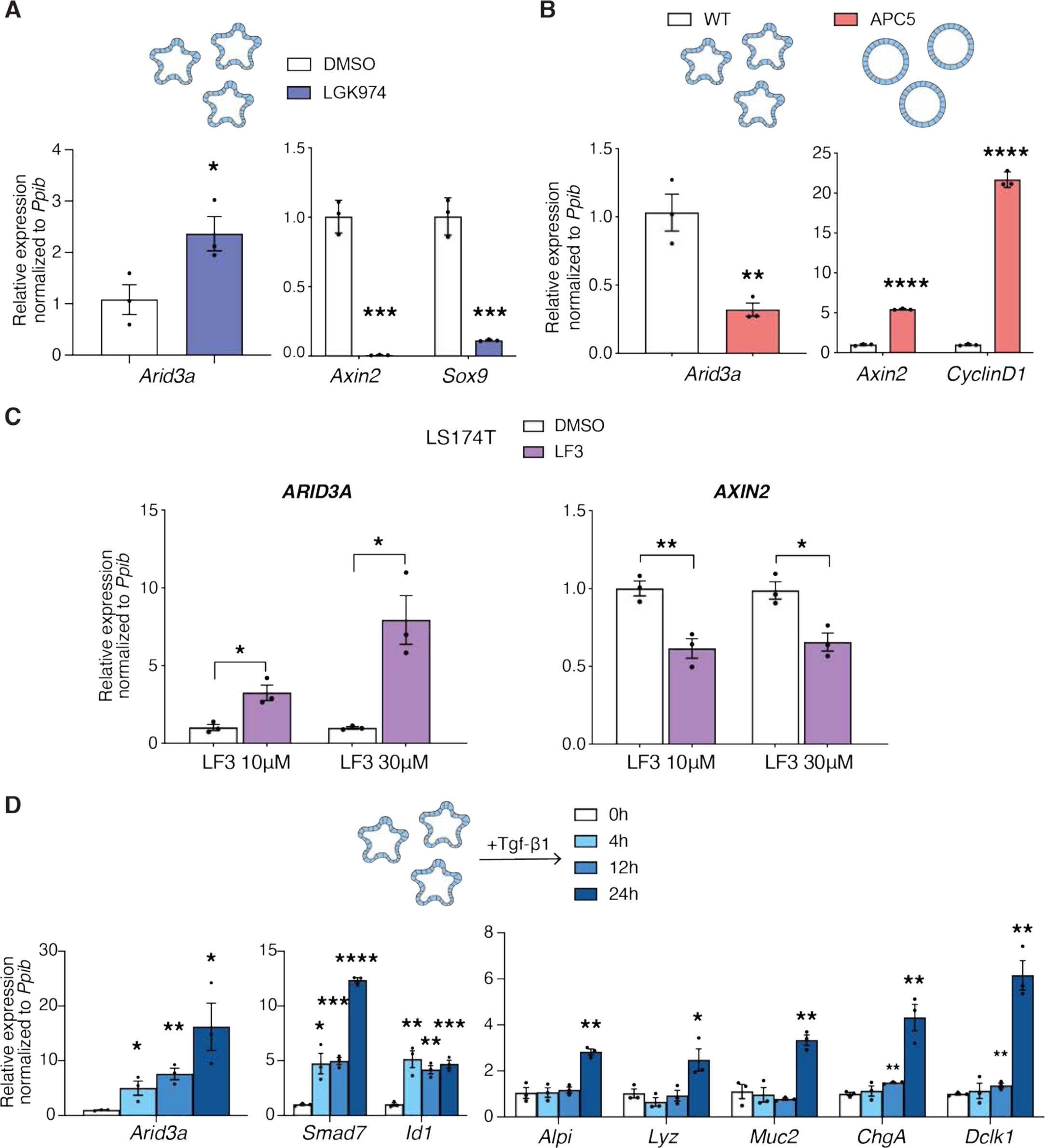
WNT and TGF-β signalling fine-tune expression of Arid3a in the small intestinal epithelium. (A) qRT-PCR analysis of WT organoids treated with LGK974 inhibitor for 48h. Organoids were established from three biologically independent animals per group (N=3). (B) qRT-PCR analysis of WT versus APC5 mutant organoids. Three independent experiments were performed (n=3). (C) qRT-PCR analysis of LS174T cells treated with two different doses of LF3. Three independent experiments were performed (n=3). (D) qRT-PCT analysis of WT organoids treated with recombinant TGF-β1 for 4h, 12h and 24h. Analysis included *Arid3a*, *TGF-β* target genes and markers of differentiation. Organoids were established from three biologically independent animals per group (N=3). Data represent mean ± s.e.m. *P<0.05, **P<0.01, ***P<0.001, two-sided t-test.

Since WNT signalling regulates ISCs self-renewal and differentiation, the expression changes of *Arid3a* upon modulation of WNT in organoids might be caused indirectly by cell fate changes. To validate if WNT directly regulates *Arid3a* without change of cell fate, we switched to the human colorectal cancer cell line LS174T - with activated WNT signalling driven by β-catenin mutation - by treating them with two different doses of LF3 inhibitor (10 and 30μΜ) for 24 hours. qRT-PCR analysis showed a robust upregulated expression of *ARID3A* and downregulation of WNT target *AXIN2* in a dose-dependent manner (Figure 2C). Immunofluorescent staining further confirmed upregulated expression of ARID3A at the protein level in LF3-treated LS174T cells (Figure S2B).

Since *Arid3a* is enriched at the early progenitor cells, we asked if its expression might be regulated by NOTCH signalling, the master regulator of early fate decision at the progenitors (7). Intestinal organoids were treated with the γ-secretase inhibitor DAPT (10μM) for 48h followed by qRT-PCR analysis. While DAPT treatment inhibited NOTCH target *Hes1* and de-repressed *Atoh1* expression, we did not observe any significant changes in *Arid3a* expression, indicating that its expression is independent of Notch signalling (Figure S2C).

Apart from WNT and NOTCH, TGF-β and BMP signalling are also involved in intestinal homeostasis, while their dysregulation has been linked to cancer and other gastrointestinal diseases (17–20). Interestingly, both TGF-β and BMP signalling form an expression gradient similar to that of *Arid3a*. To test whether the expression of *Arid3a* is regulated by the TGF-β superfamily, we treated mouse intestinal organoids with recombinant TGF-β1 and BMP4 ligands. As expected, BMP treatment activated its target gene *Id1* and inhibited ISC marker *Lgr5* expression (21, 22). However, the expression of *Arid3a* was unaffected by BMP (Figure S2D). In contrast, treatment of organoids with recombinant TGF-β1 for 4h, 12h and 24h revealed that *Arid3a* expression was increased in a dose-dependent manner, similar to the expression of TGF-β target genes *Smad7* and *Id1* (Figure 2D). It is important to note that the intestinal differentiation markers were not upregulated until 24h after TGF-β induction, indicating that *Arid3a* expression was driven by TGF-β signalling directly rather than because of increased differentiation (Figure 2D).

### *Arid3a* regulates transit-amplifying cell proliferation

It has been previously reported that *Arid3a*-null mice exhibit embryonic lethality due to defects in haematopoiesis (23). In order to characterise the functional role of ARID3A in intestinal epithelial cells, we generated an *in vivo* conditional knockout mouse model, *Villin*Cre-ERT2+/-; *Arid3a*fl/fl (*Arid3a* cKO), to delete *Arid3a* in *Villin*+ intestinal epithelial cells upon tamoxifen induction (24, 25). Analysis of intestinal tissues 1 month post-tamoxifen administration confirmed complete abolishment of ARID3A expression at the epithelial cells, while the expression remained unchanged at stromal cells (Figure 3A). Haematoxylin and Eosin (H&E) staining did not show any noticeable changes in the gross morphology of the *Arid3a*-depleted intestine (Figure S3A). However, semi-quantitative pathologist scoring revealed that *Arid3a* cKO animals exhibited a minimal to mild villus atrophy (Figures S3A and S3B). Moreover, *Arid3a* cKO mice were also found to have shorter small intestine (mean= 35.35cm) when compared to WT (mean= 37.60cm) (Figure 3B).

**Figure 3.**
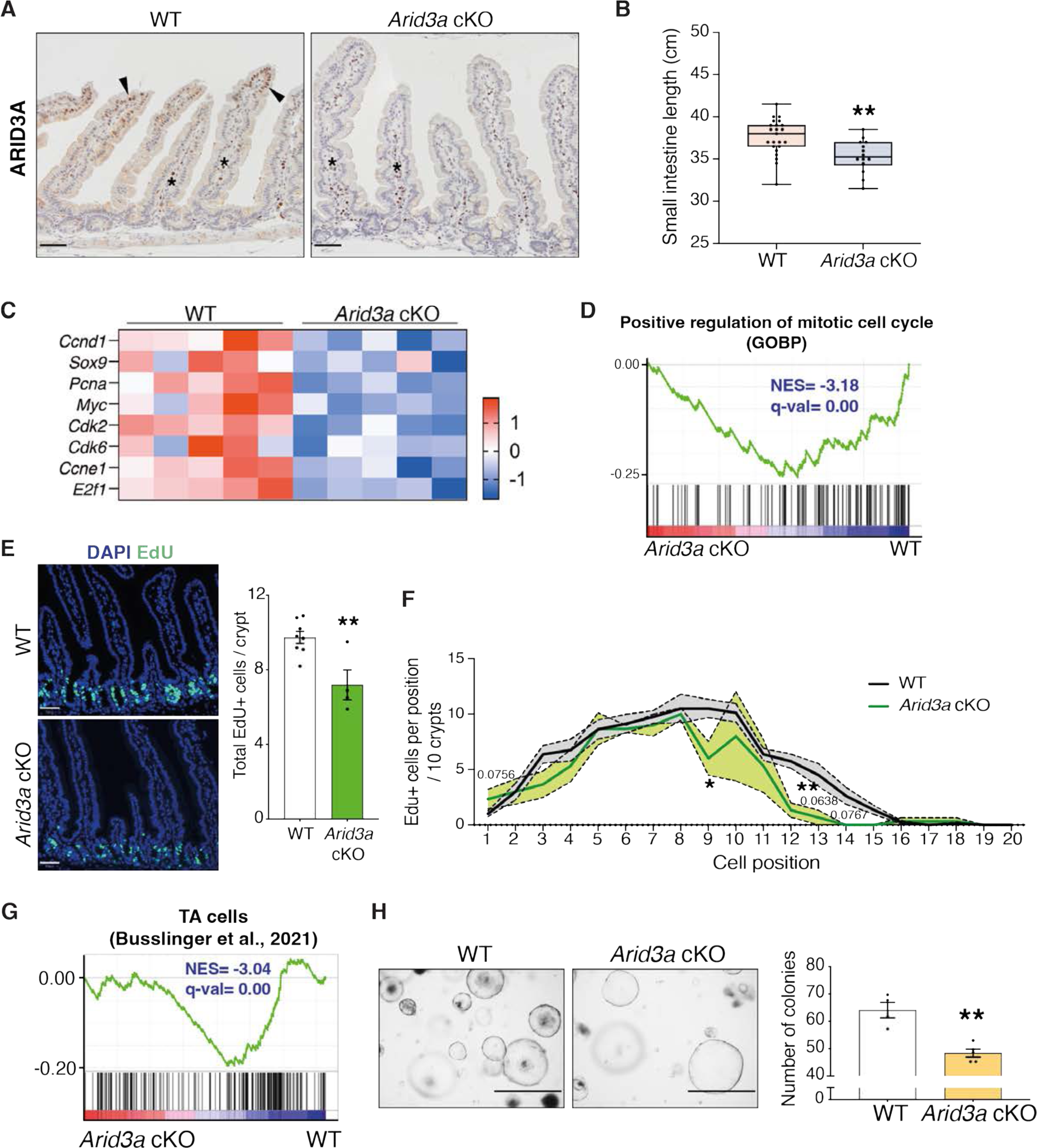
Loss of *Arid3a* leads to loss of proliferative TA cells. (A) ARID3A immunostaining of WT and *Arid3a* cKO mice. Representative images N=3 of each group. Scale bar, 50μm. (B) Differences small intestinal length (cm) at 1 month after tamoxifen administration. N=23 WT animals and N=14 cKO animals. Box plot shows all datapoints from min to max. *P<0.05, **P<0.01, ***P<0.001, two-sided t-test. (C) Heatmap of RNA-seq data of representative proliferation markers. Z-scores are shown. (D) GSEA of GO biological process “Positive regulation of mitotic cell cycle”. (E) EdU staining of WT and *Arid3a* cKO animals. Representative picture of N=8 WT and N=4 cKO animals. Scale bar, 50μm. Quantification of total EdU positive cells per crypt. Data represent mean ± s.e.m. *P<0.05, **P<0.01, ***P<0.001, two-sided t-test (F) Quantification of EdU-positive cells based on their crypt position (per 10 crypts). Data represent mean ± s.e.m. (light colour area) *P<0.05, **P<0.01, ***P<0.001, multiple two-sided t-tests. (G) GSEA of previously published TA cells gene lists. (H) Images of WT and *Arid3a* cKO organoid formation assay and quantification of organoids per genotype at Day 5 after isolation. Representative picture of N=4 WT and N=5 animals. Scale bar, 1000μm. Data represent mean ± s.e.m. *P<0.05, **P<0.01, ***P<0.001, two-sided t-test.

To characterise the molecular changes caused by *Arid3a* deletion, unbiased RNA sequencing (RNA-seq) analysis was performed on the *Arid3a* cKO and WT intestine. Hierarchical clustering analysis showed that samples with the same genotypes readily clustered together (Figure S3C). Differential gene expression analysis revealed a total of 4387 genes differentially expressed between the two groups (FDR cut-off <0.05), with 2413 genes upregulated and 1974 downregulated in the *Arid3a* cKO intestine (Supplemental information 1). In particular, we observed a mild to moderate decrease of various cell cycle and cell proliferation markers in the *Arid3a*-depleted intestinal crypts (Figure 3C). Indeed, Gene Set Enrichment Analysis (GSEA) of mitotic cell cycle regulators indicated a strong downregulation in cKO intestine (Figure 3D), while Metacore analysis identified the regulation of cell population proliferation as the most significantly affected Gene Ontology (GO) biological process (Figure S3D). To assess the number of mitotically active cells in the crypts, animals were injected with a short pulse of 5-ehtynyl-2’deoxyuridine (EdU) to label cells undergoing *de novo* DNA synthesis or S-phase synthesis of the cell cycle. In accordance with our previous findings, *Arid3a* cKO intestine showed a reduction in total numbers of proliferative cells per crypt compared to WT animals (WT mean=9.73 cells; cKO mean=7.2 cells) (Figure 3E). Interestingly, quantitation of EdU-positive cells revealed that reduced proliferation was mostly observed at cell positions 9-15 (counting from the crypt bottom), whereas EdU-labelling was largely unchanged at cell positions 1-5 (Figure 3F). This result suggests that *Arid3a* depletion inhibits proliferation at TA cells in the upper crypt whilst ISCs at the crypt base are mostly unaffected. Indeed, GSEA revealed a significant transcriptional downregulation of and TA cell gene signature (26) as well as in WNT signalling (27) at *Arid3a* cKO crypts (Figures 3G and S3E). Of note, RNAscope analysis confirmed no difference in the expression of the ISC-specific marker *Olfm4* between WT and *Arid3a* cKO tissues, indicating that the number of stem cells are not affected upon *Arid3a* deletion (Figure S3F).

To validate the *in vivo* data, we generated *ex vivo* organoid cultures for functional analysis. Organoid formation analysis showed a significant decrease in organoid formation capacity in *Arid3a* cKO derived crypts (WT mean=64.1 organoids, cKO mean=48.4 organoids) (Figure 3H). We further challenged the organoids by depleting one of the essential growth factors and WNT agonist, RSPONDIN (RSPO), in the culture medium to test the organoid dependency on exogenous WNT signal. Murine organoid cultures rely on supplementation of exogenous growth factors RSPO, EGF and NOGGIN to survive (28). Under normal condition, organoids were cultured in medium containing 5% RSPO conditioned media (CM). Organoids derived from WT and cKO animals were assessed and quantified based on their morphologies: organoids with >3 buds were considered healthy, organoids with 1-3 buds were considered unhealthy and organoids that failed to bud or form cysts were considered collapsed. WT and cKO organoids did not exhibit any major morphological differences in the presence of 5% RSPO CM (Figure S3G). However, when organoids were challenged with a lower RSPO concentration (1%), a higher percentage of collapsed and a lower percentage of healthy organoids were observed (Figure S3G). The increased dependence on exogenous RSPO in the cKO organoids suggests a reduction of endogenous WNT signalling in the *Arid3a*-depleted crypt cells, which is consistent with the GSEA data observed in Figure S3E.

### Loss of *Arid3a* perturbs absorptive and secretory cell differentiation

Next, we asked if functional differentiation is affected in the cKO intestine. We first looked into differentiation of Paneth cells, the only specialised epithelial cells that reside at the crypt base adjacent to ISCs (7). RNA-seq analysis revealed downregulation of various Paneth cell markers in *Arid3a* cKO intestine, including *Lyz1*, the newly discovered marker *Mptx2* (29), as well as a large number of anti-microbial peptides (α-defensins, *Defa*) secreted specifically from Paneth cells (30) (Figure 4A). Immunostaining of LYZ further confirmed the reduction of Paneth cells numbers in the cKO intestine (WT mean=14.53 cells/5 crypts, cKO mean=11.82 cells/5 crypts) (Figure 4B). Paneth cells function as ISC niche by secretion of essential niche factors such as WNT ligands (31). Reduced Paneth cell numbers in Arid3a-deficient intestine may shorten the WNT signal gradient in the crypt, leading to the observation of reduced WNT signature and TA cell proliferation.

**Figure 4.**
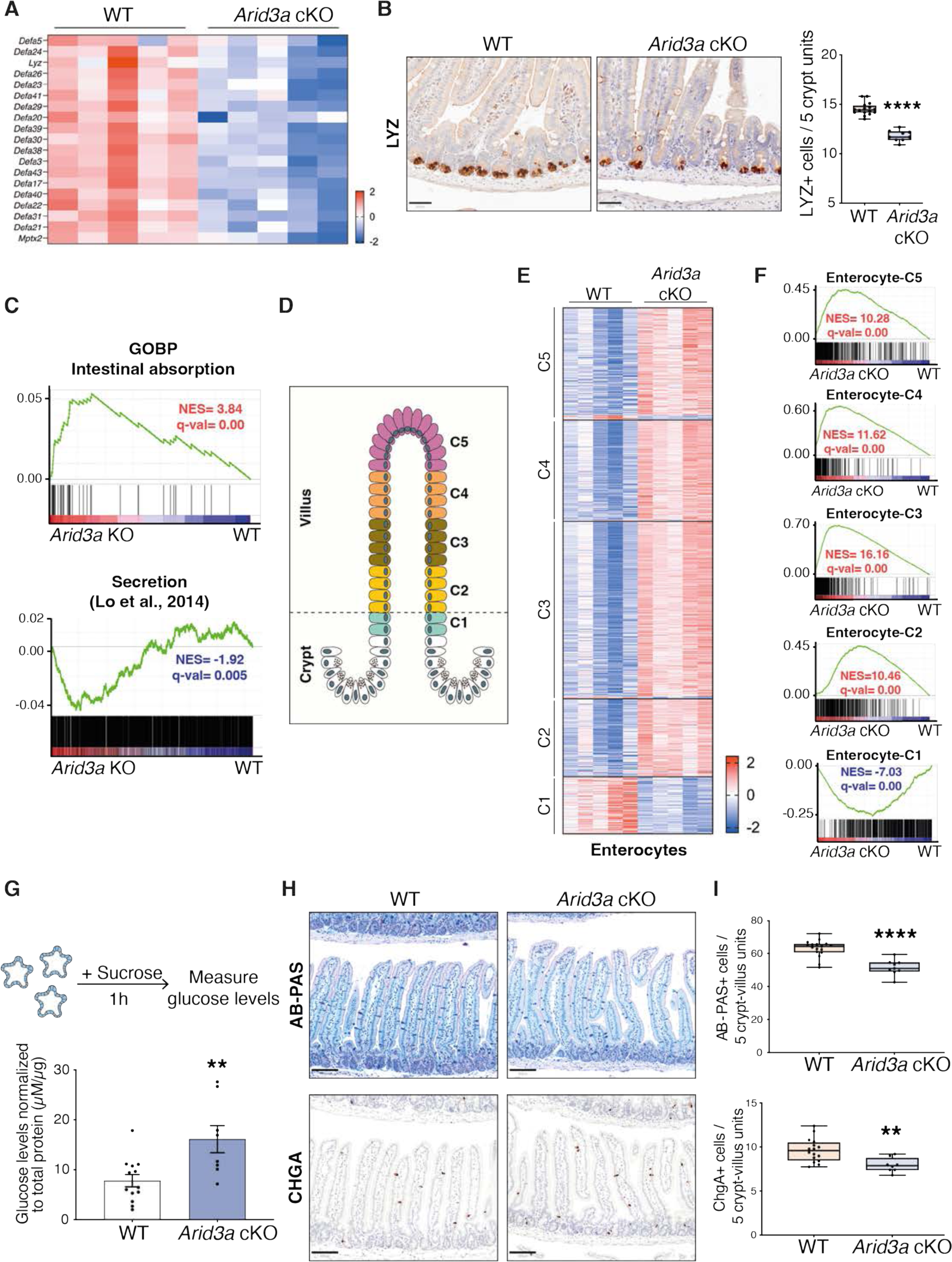
Loss of Arid3a leads to upregulation of mid-villus and villus tip gene signatures of all lineages. (A) Heatmap of RNA-seq data of representative Paneth cells markers. Z-scores are shown. (B) LYZ staining of WT and Arid3a cKO animals. Scale bar, 50 μm. Representative images and quantification of N=16 WT and N=9 KO animals. Box plot shows all datapoints from min to max. *P<0.05, **P<0.01, ***P<0.001, two-sided t-test. (C) GSEA of intestinal absorption and secretion related genes. (D) Illustration of enterocyte zonation model as proposed by Moor et al., 2018. (E) Heatmap of differentially expressed genes of all enterocyte clusters based on RNA-seq data. Z-scores are shown. FDR cut-off <0.05. (F) GSEA of all genes included in the 5 enterocyte clusters (Moor et al., 2018). (G) Disaccharide assay: WT and *Arid3a* cKO organoids were treated with sucrose for 1h. Graph shows absorbance levels of glucose normalised to total protein. Data represent mean ± s.e.m. *P<0.05, **P<0.01, ***P<0.001, two-sided t-test. (H) AB-PAS and CHGA staining of WT and Arid3a cKO animals. Scale bar, 100μm. Representative images of N=17 WT and N=9 KO animals for AB-PAS and N=16 WT and N=8 KO animals for CHGA. (I) Quantification of AB-PAS and CHGA stainings. Box plot shows all datapoints from min to max. *P<0.05, **P<0.01, ***P<0.001, two-sided t-test.

In adult intestinal crypts, ISCs give rise to progenitor cells at the TA zone that will adopt either absorptive or secretory fate. Apart from Paneth cells, all other differentiated epithelial cells migrate up towards the villus after lineage commitment. GSEA showed significant enrichment of absorptive gene signature and an overall reduced secretion signature in the mutant intestine (Figure 4C), suggesting that Arid3a regulates the differentiation ratio between absorptive and secretory cells. Recent studies further revealed that the differentiated cells do not acquire their terminal identity at the TA zone. Rather, the committed cells are further zonated along the villi to carry out different functions (10, 11). In particular, enterocyte zonation can be grouped into five distinct functional clusters across the crypt/villus axis (11) (Figure 4D). Interestingly, differential gene expression analysis and GSEA showed that Cluster 1 (early crypt enterocytes) was strongly enriched at WT animals, while the gene signatures of Clusters 2-5 were upregulated in *Arid3a* cKO intestines (Figures 4E and 4F). Immunostaining of the villus tip-enriched markers APOA4 and ASPA confirmed their upregulation and perturbed zonated expression in the cKO intestines (Figures S4A and S4B). It has been previously shown that genes in Cluster 2 are associated with mitochondrial activity and clusters 3-5 are associated with absorption, nutrient transport, brush border function and cell adhesion (11). Interestingly, Metacore analysis of the significantly upregulated genes of the RNA-seq dataset (FDR<0.05, Fold change>1.5) showed that most of the top upregulated GO biological processes were related to an increase of different metabolic processes (Figure S4C), which are likely facilitated by the increased enterocyte absorptive gene signatures in Clusters 2-5. Furthermore, disaccharidase functional assay confirmed an increased sucrose breakdown to glucose in the *Arid3a* cKO organoids, indicating higher enterocyte digestion activity in the *Arid3a*-depleted intestinal epithelium compared to WT control (Figure 4G).

Contrary to the enterocyte differentiation, secretory lineage appeared to be downregulated in Arid3a-deficient intestine (Figure 4C). Apart from Paneth cells, goblet and enteroendocrine cells are the two most abundant secretory cell types in the intestinal epithelium throughout the villus. Indeed, staining of Alcian Blue / Periodic Acid-Schiff (AB-PAS) and Chromogranin A (CHGA) showed minimal reduction of goblet and enteroendocrine cells, respectively, in the *Arid3a* cKO intestine (Figure 4H). Similar to enterocytes, several recent studies have reported the spatial differentiation dynamics of secretory cells (10, 32, 33). Goblet cells and tuft cells show a spatial expression programme across the crypt-villus axis similar to the one of enterocytes, while enteroendocrine cells show a more complex spatio-temporal migration pattern (10, 33). Interestingly, despite of the observed overall reduced secretory cell differentiation, we noted a subtle but significant enrichment of the villus-tip gene expression programme across all three secretory lineages upon *Arid3a* deletion (Figures S4D-I), suggesting that ARID3A may regulate terminal differentiation of secretory cells at the upper villus. Together, our data indicates that deletion of ARID3A impairs enterocytes versus secretory cell differentiation ratio and spatial gene signatures of the intestinal epithelium.

### Single-cell RNA-sequencing reveals changes in enterocyte differentiation trajectory and cell cycle phase in TA cells

Our bulk transcriptomics analysis in combination with tissue analysis revealed changes in TA cell proliferation and cell type composition in the *Arid3a* cKO intestine. To better understand how ARID3A regulates the dynamics of cell proliferation and differentiation, we further performed single-cell RNA-sequencing (scRNA-seq) on the intestinal epithelium from the WT and mutant animals. Clustering of epithelial cells from both WT and *Arid3a* cKO intestine showed 12 distinct clusters including all the known epithelial cell types and curated cell type annotation was performed based on previously published markers of each cell population (Figure S5A, Supplemental Information 2). UMAP analysis further revealed 5 distinct enterocytes clusters that exhibit similarities with the previously described zonated clusters along the villus axis (11) (Figure 5A). In accordance with the results observed earlier, we found that loss of *Arid3a* resulted in an increase in enterocyte populations (WT=51.3%, cKO=56%) and a reduction of secretory cells (WT=15.8%, cKO=12.4%) and TA cells (WT=13.2%, cKO=12.3%), whilst the number of Lgr5+ ISCs remains largely unaffected (Figure 5B).

**Figure 5.**
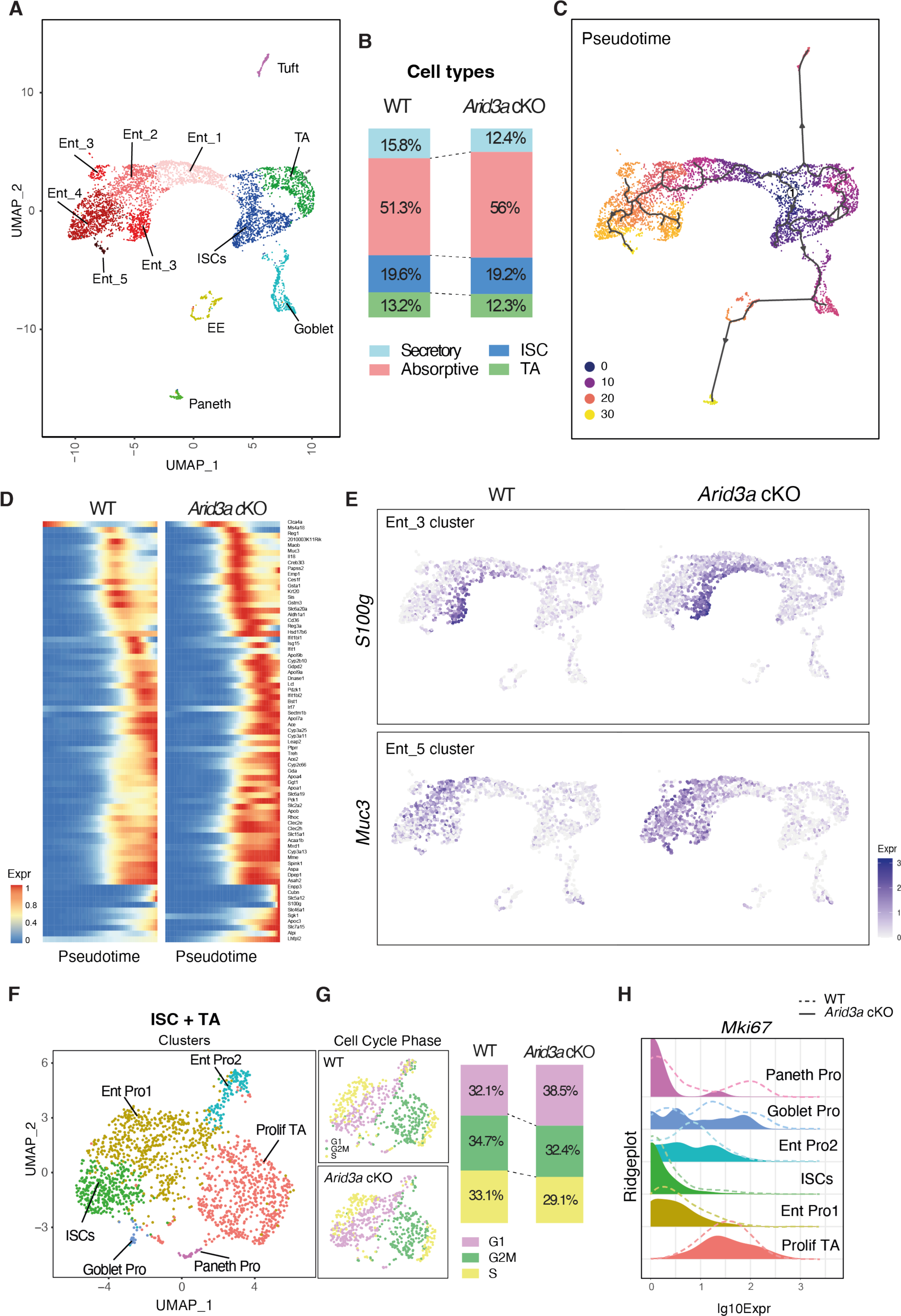
Single-cell RNA-sequencing reveals changes in enterocyte differentiation trajectory and cell cycle phase in TA cells. (A) UMAP plot of epithelial cells from WT and *Arid3a* cKO intestine and post-hoc annotation of different cell types. (B) Proportion of intestinal cell types in WT and *Arid3a* cKO animals based on scRNA-seq. (C) Pseudotime analysis of the scRNA-seq data. ISCs were chosen as the starting point of the trajectory analysis. (D) Array of gene expression over pseudotime of the most DEG that are significant in the enterocyte clusters based on RNA-seq data.(E) Feature plot of *S100g* (enterocyte marker cluster 3) and *Muc3* (enterocyte marker cluster 4) expression on UMAP. (F) UMAP plot of ISC+TA cells from WT and *Arid3a* cKO intestine and post-hoc annotation of different sub-populations within this compartment. (G) Feature plot of cell cycle phase on UMAP of the ISC+TA compartment of WT and *Arid3a* cKO. On the right, proportion in percentages of cells in G1, G2/M or S phase in the WT and *Arid3a* cKO intestines.(H) Ridgeplot showing log10 expression of *Mki67* gene in WT and *Arid3a* cKO for each crypt cell sub-population.

To better understand the changes in differentiation and cell fate dynamics, we performed pseudotime trajectory analysis which revealed 3 distinct major trajectories: differentiation trajectory towards 1) enterocyte lineage, 2) Goblet, EE and Paneth cells and 3) Tuft cells (Figure 5C). This indicates that Tuft cells is a distinct cell lineage from other secretory cell types. Loss of *Arid3a* did not cause major alteration to the overall differentiation dynamics. Since we observed a significant increase in enterocyte signatures in *Arid3a* cKO intestine from bulk RNA-seq data, we ask if loss of *Arid3a* would affect enterocyte differentiation dynamics. Indeed, spatial mapping of the most differentially expressed enterocyte genes (based on the bulk RNA-seq) against pseudotime showed enriched expression of these genes towards earlier pseudo-timeline in the absence of *Arid3a* (Figure 5D). This was confirmed by mapping the expression of selected enterocyte markers (*S100g, Muc3, Ms4a18* and *Ace2*) in UMAP, which showed increased expression of these genes in the *Arid3a* cKO cells aligned at earlier pseudo-temporal trajectory compared to WT (Figure 5E and Figure S5B), suggesting a role of *Arid3a* in enterocyte differentiation dynamics.

We have previously shown reduced crypt proliferation in *Arid3a* cKO intestine. To further understand the proliferative changes in different crypt cell populations, ISCs and TA cell clusters were subjected to additional sub-clustering analysis. This resulted in 6 cell clusters, including ISCs, proliferative TA cells, Paneth progenitors, Goblet progenitors and two Enterocyte progenitor populations (Figure 5F). Cell cycle analysis of these cells showed an overall increase in G1 phase (WT=32.1%, cKO=38.5%) and a corresponding decrease in G2M (WT=34.7%, cKO=32.4%) and S phase (WT=33.1%, cKO=29.1%) in the mutant crypt cells (Figure 5G). Consistent with the EdU quantitation result observed earlier, we noted reduced expression of proliferation markers (*Mki67*, *Pcna* and *Ccna2*) in different crypt cell populations of the mutant animals, notably more prominent in the proliferative TA cells and less affected in the ISCs (Figure 5H and Figure S5C).

Together, our data shows that *Arid3a* deletion resulted in reduced proliferation in the TA cells and enhanced enterocyte differentiation. Transcriptomic analysis using both bulk and scRNA-seq indicates that ARID3A modulates intestinal differentiation dynamics in the villus by fine-tuning the expression level of the spatial differentiation markers at the corresponding zones.

### Increased binding and transcription of HNF1 and HNF4 in *Arid3a*-depleted intestine

In order to understand how loss of *Arid3a* perturbs intestinal spatial differentiation and causes changes in cell composition of the small intestinal epithelium, we performed ATAC-seq to assess the differences in chromatin accessibility between WT and *Arid3a* cKO intestine. Analysis of the ATAC-seq data showed that the *Arid3a* cKO intestine exhibits a more global open chromatin pattern when compared to WT (Figure S6A). Although these changes were mild, the result was unexpected considering that *Arid3a* deletion caused increased differentiation that is often associated with less open chromatin (34). This suggests that the overall enhanced chromatin accessibility may not fully explain the perturbation of proliferation and differentiation caused by *Arid3a* loss. To gain more insight into the mechanism of ARID3A function, we performed footprinting analysis of our ATAC-seq using a recently published methodology – Transcription factor Occupancy prediction By Investigation of ATAC-seq Signal (TOBIAS) (35), which enables genome-wide analysis of transcription factor (TF) dynamics and calculates enriched motif binding using publicly available binding motifs of hundreds of transcription factors. Interestingly, TOBIAS analysis of the ATAC-seq data showed an enrichment of transcription factor binding sites in AT-rich genomic regions of the Arid3a cKO intestine (Figure 6A). These included members of the ARID family (ARID3B and ARID5A) as well as members of the HNF family (Hnf1 and Hnf4). On the other side, WT intestine showed enrichment of binding motifs of FOS/JUN dimers (AP-1 pathway) (Figure 6A), which has been previously linked with proliferation, apoptosis and carcinogenesis (36). To confirm this result, we utilised ChromVAR to assess the TF-associated chromatin accessibility. TFs with variability score >5 are associated with dynamic chromatin contributing to phenotypic changes, while TFs with variability score <5 are associated with permissive chromatin (37). In accordance with the TOBIAS analysis, FOS/JUN dimers and HNF4A-associated motifs were at the top two highest variability scores (Figure 6B), indicating that changes of these TF dynamics contribute to the altered proliferation and differentiation in *Arid3a* cKO intestine.

**Figure 6.**
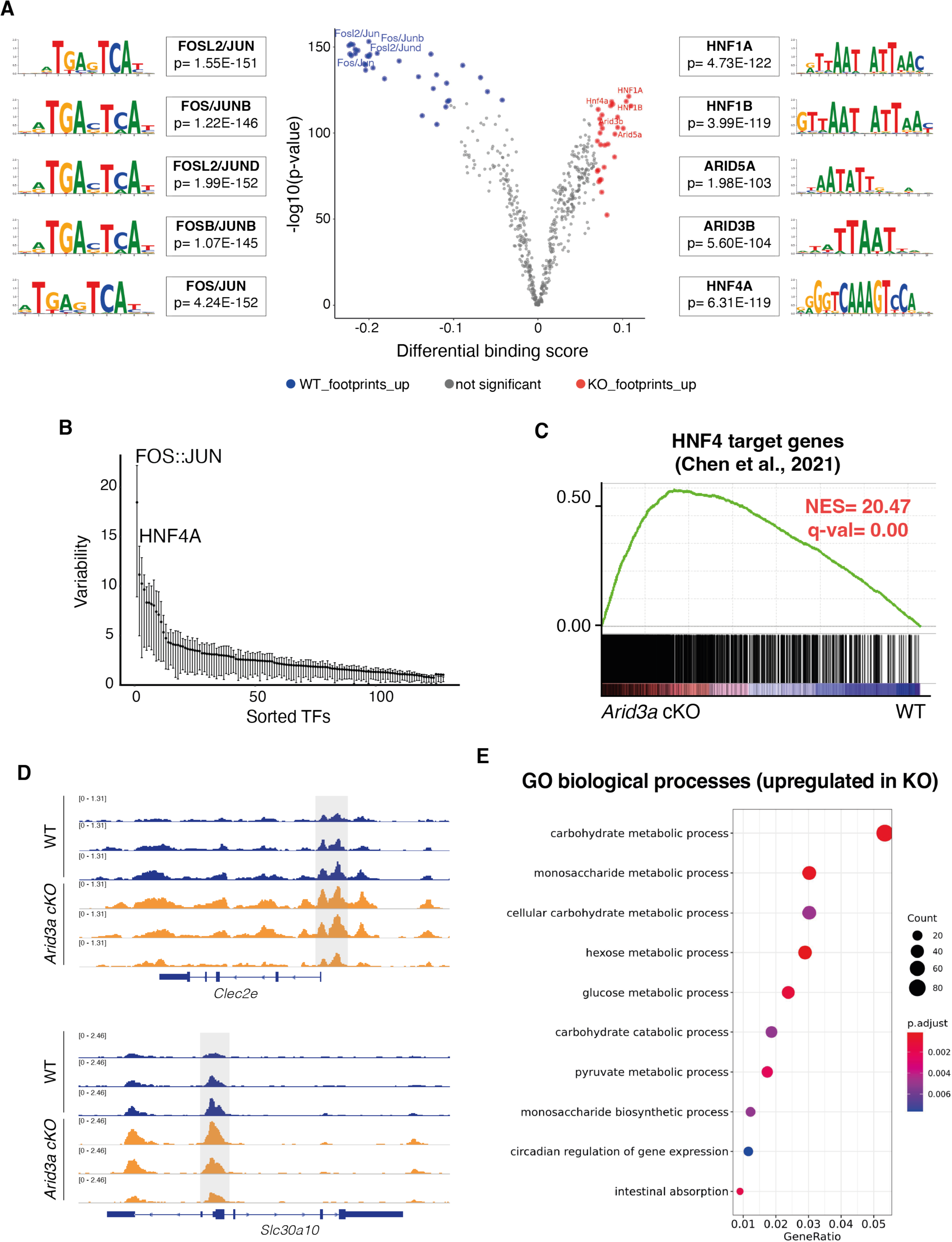
Deletion of Arid3a allows HNF-family of transcription factors to bind to A+T rich regions. (A) Analysis of the ATAC-seq using the TOBIAS package (see methods). Volcano plot shows the differential binding activity against the -log10(p-value) of all investigated transcription factor motifs. Each dot represents one motif; blue dots represent motif enrichment in WT; red dots represent motif enrichment in *Arid3a* cKO. Representative examples of Top-10 transcription factors enriched in either WT or *Arid3a* cKO animals is shown of the left and right side of the volcano plot, respectively. (B) Analysis of the ATAC-seq using the ChromVAR package (see methods). Plot shows TFs sorted based on their variability score. (C) GSEA of previously published gene list of HNF4 target genes. (D) Comparison of open ATAC-seq chromatin peaks of Hnf4 targets genes (*Clec2e* and Slc30a10) in WT and *Arid3a* cKO animals (tracks extracted from IGV). (E) Top 20 upregulated GO biological processes based on ATAC-seq analysis. GO was performed based on 5000 peaks with the greatest variability between WT and KO.

Interestingly, HNF proteins have been previously associated with terminal maturation of enterocytes (38–40). It is conceivable that enrichment of HNF activity in the *Arid3a* cKO intestine results in increased spatial differentiation of enterocytes along the villus. Indeed, GSEA of the RNA-seq data showed a strong enrichment of the previously published Hnf4a/g target genes in the *Arid3a* cKO intestine (Figure 6C), confirming the increased HNF4 transcription in the absence of *Arid3a*. In accordance with this finding, previously described intestinal epithelial-specific HNF4 targets (*Clec2e* and *Slc30a10*), showed higher chromatin accessibility in the promoter regions of the *Arid3a*-deficient intestine compared to WT (Figure 6D). We further performed GO analysis of the 5000 most variable peaks between WT and *Arid3a* cKO intestine to evaluate the biological processes behind. Interestingly, we observed significant upregulation of many processes related to enterocyte functions in the cKO intestine including carbohydrate/monosaccharide metabolic processes and intestinal absorption (Figure 6E), supporting the notion that HNF-mediated terminal differentiation of enterocytes is enriched in the mutant. Together with the transcriptomic data, we propose that ARID3A functions to maintain cell proliferation and prevent pre-mature HNF-mediated terminal differentiation at the TA progenitor cells.

### Loss of *Arid3a* impairs irradiation-induced regeneration

Since deletion of *Arid3a* inhibits proliferation of the TA cells where progenitors reside, we asked whether this would affect the regenerative capacity of the intestine upon irradiation. Intestinal epithelium regenerates rapidly within days after irradiation: (1) apoptotic phase (1-2 days post-irradiation, dpi), (2) hyperproliferating/regenerating phase (3-4 dpi) and (3) normalisation phase (5 dpi onwards) (41). We have shown earlier that *Arid3a* forms an expression gradient from villus tip to the early progenitor cells at the crypt (Figures 1C and 1D). We first asked if ARID3A expression is changed upon irradiation. To address that, we irradiated WT mice and collected the irradiated intestinal tissues at 1, 2, 3 and 4 dpi as well as the non-irradiated controls. Immunohistochemistry analysis showed collapsed crypts and transient loss of *Arid3a*+ cells on 1dpi, followed by crypt elongation and increased numbers of *Arid3a*+ cells at the upper crypt during the regenerating phase (3dpi). Approaching the normalisation phase on day 4, the number of ARID3A+ cells returned to homeostatic levels (Figure S7A). This suggests that *Arid3a* may play a role in intestinal regeneration and restoration of tissue homeostasis after injury.

Next, we investigated whether deletion of *Arid3a* perturbs the regenerative response. Expert pathological analysis was performed on the irradiated tissues to assess the damage of the lamina propria, mucosa and gut-associated lymphoid tissue, and confirmed a more extensive tissue damage of cKO animals from day 2 to day 4 when compared to WT (Figure S7B and S7C). We then performed immunostaining of KI67 and Cleaved caspase-3 (c-CASP3) to assess tissue proliferation and apoptosis respectively (Figure 7A and 7B). No major differences were observed between WT and cKO animals on 1 dpi, where extended cell death led to a dramatic reduction of proliferation. On 2 and 3 dpi, WT intestine started regenerating as evident by crypt expansion and increased proliferation, whilst *Arid3a* cKO intestine showed much lower numbers of KI67+ cells with minimal crypt expansion. By 5 dpi, WT intestine had returned to the normalisation phase, whereas the majority of *Arid3a* cKO crypts were still in the hyperproliferation phase with elongated crypts (Figure 7A). To confirm the reduced proliferative capacity of crypts at early timepoints, we isolated crypts from WT and *Arid3a* cKO intestine collected on 1 and 3 dpi and performed organoid formation assay. No differences were observed on day 1 apoptotic phase where organoid formation efficiency was less than 10% (Figure S7D). However, on 3 dpi, WT crypts had restored their capacity to form organoids while the colony formation efficiency remained low for the *Arid3a* depleted crypts (p-value=0.099) (Figure S7D). This is consistent with the earlier observation that the regeneration phase of *Arid3a* cKO intestine was impaired on 3 dpi.

**Figure 7.**
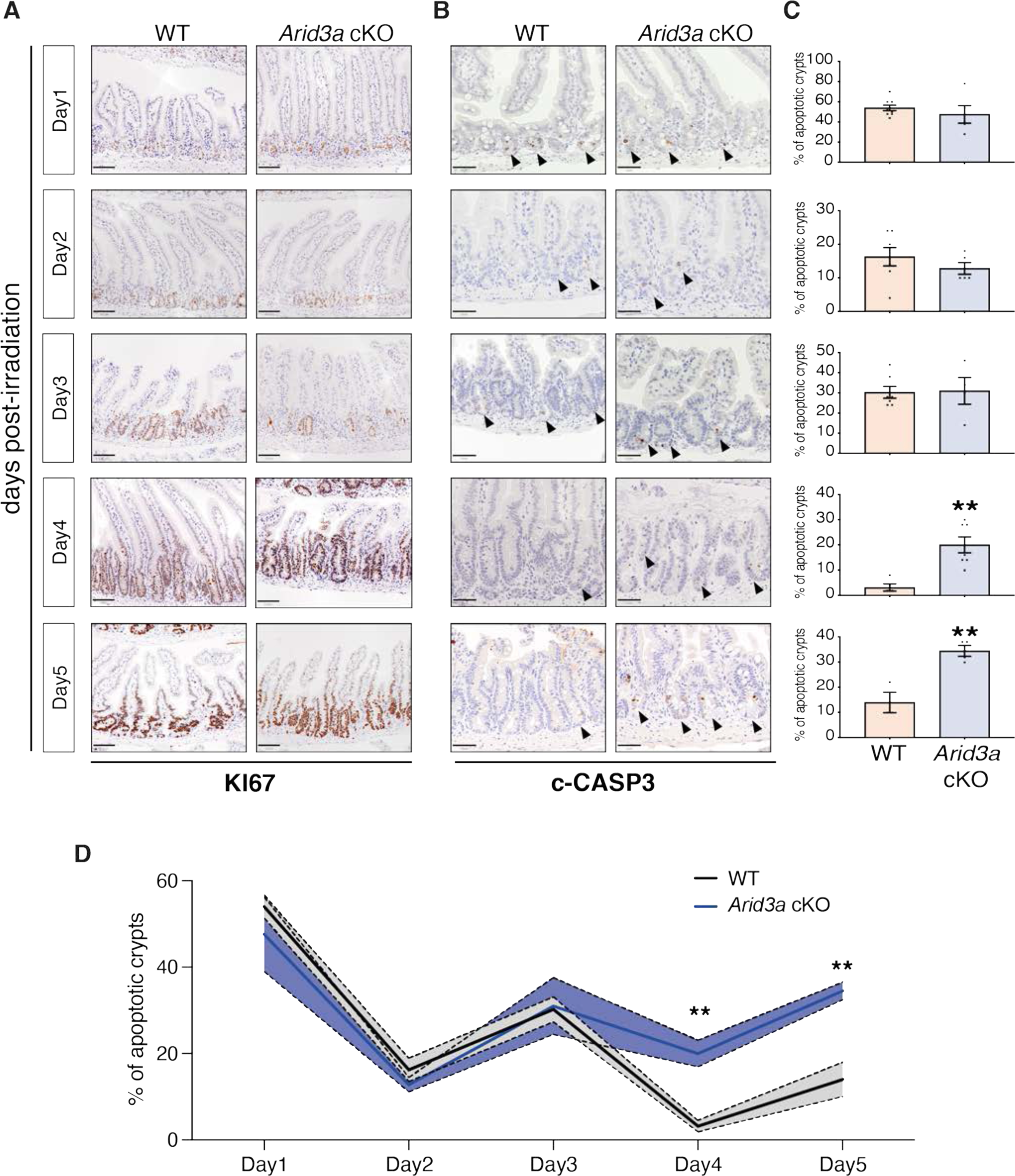
*Arid3a* cKO animals exhibit a delayed irradiation-induced regeneration. (A) KI67 immunostaining of WT and *Arid3a* cKO mice. Representative images of at least N=4 animals for each genotype per timepoint. Scale bar, 100μm. (B) c-CASP3 immunostaining of WT and *Arid3a* cKO mice. Representative images of at least N=4 animals for each genotype per timepoint. Scale bar, 50μm. Black arrowheads indicate apoptotic cells. (C) Quantification of apoptotic crypts for each timepoint is shown on the right side. Data represent mean ± s.e.m. *P<0.05, **P<0.01, ***P<0.001, two-sided t-test. (D) Summary of quantification of apoptotic crypts in WT and Arid3a cKO animals shown in (C). Data represent mean ± s.e.m. (light colour area) *P<0.05, **P<0.01, ***P<0.001, two-sided t-test.

Interestingly, c-CASP3 staining showed no differences in the number of apoptotic cells between WT and mutant at the initial apoptotic and regenerating phase (1-3 dpi) (Figures 7B and 7C), suggesting that perturbation of regeneration was not directly caused by increased apoptosis. However, approaching the normalisation phase (4-5 dpi), *Arid3a* cKO intestine showed a significantly higher percentage of apoptotic crypts compared to control (Figures 7B and 7C). It is interesting to note that the number of apoptotic crypts in both WT and mutant intestine appeared to decrease gradually over time in a wave rather than linear pattern (Figure 7D), suggesting that crypts unable to regenerate in the first wave after irradiation may collapse again. On the other hand, the apoptotic wave in the *Arid3a*-depleted intestine failed to subside and remained high over time (Figure 7D). Of note, increased expression of apoptotic markers was also observed in the cKO intestine during homeostasis, suggesting that ARID3A may also play a role in regulating apoptosis (Figure S7E).

## Discussion

ARID3A is a transcription factor with DNA binding domain that interacts with A+T rich genomic regions (42, 43). Functionally, ARID3A has been shown to drive normal development of both myeloid and B cell lineage specification, whereas deletion of the mouse *Arid3a*, leads to embryonic lethality due to defective haematopoiesis (44). Interestingly, *Arid3a* also regulates B cell response to antigen via post-translational palmitoylation of cytoplasmic ARID3A, leading to lipid rafts accumulation and B-cell antigen receptor (BCR) signalling (45). Moreover, ARID3A is enriched in megakaryocytes compared to haematopoietic progenitor cells and has been shown to promote terminal megakaryocytic differentiation (46). However, the role of ARID3A in intestinal epithelium has not yet been explored. A recent single-cell analysis of the developing gut has identified *Arid3a* as one of the key regulators of intestinal epithelial development through transcription factor regulatory network mapping, whilst the mechanism remains unknown (47).

Here, we report for a previously unrecognised role of ARID3A in adult intestinal epithelial differentiation. We showed that ARID3A is regulated by WNT and TGF-β signalling to generate an expression gradient from villus tip to TA cells at the crypt. Loss of *Arid3a* in the intestinal epithelium leads to reduced TA cell proliferation and increased enterocyte differentiation, suggesting that ARID3A plays a critical role in coordinating the proliferation-differentiation switch of the TA zone in the intestine. Expression of ARID3A in the villi may also contribute to the fine-tuning of the spatial differentiation programme along the villus by regulating their expression level. Increased binding and transcription of the downstream targets of HNF4 in the mutant intestine suggests that ARID3A may function to maintain TA cell proliferation by inhibiting HNF4-mediated differentiation. We further showed that reduced TA cell proliferation in the *Arid3a* cKO intestine impairs intestinal regenerative potential.

It is well understood that TA cells in the intestinal crypts contain progenitors that will continue to proliferate, fuelling the intestinal epithelium in the villus, whilst undergoing differentiation in parallel to adopt one of the functional cell types (48). However, the molecular control of proliferation-differentiation switch in TA cells has not been formally characterised. Previous studies of the intestinal epithelium focused mostly on intestinal stem cells and the NOTCH-dependent binary fate decision at the early +4/5 cell progenitors, while regulation of TA cells is largely overlooked. A recent study showed that modulation of TA cell proliferation changes the balance of absorptive to secretory cell ratio (49), further highlighting the central role of TA cell regulation in intestinal homeostasis. Our current findings provide a new cell intrinsic regulatory mechanism of TA cells, where ARID3A functions to coordinate the proliferation-differentiation switch by regulating HNF4 binding and its target gene transcription.

Besides the binary cell fate decision at the early progenitors, it has been recently shown that both absorptive and secretory cells undergo spatial differentiation along the villus axis, resulting in regional and functional heterogeneity of all cell types (10, 11). This implies that intestinal differentiation is a continuous process throughout the crypt-villus axis. More recent studies have shown that the zonation patterning of enterocytes is regulated by a BMP signalling gradient or villus tip telocytes (12, 50), while BMP’s downstream target c-MAF has also been reported to act as a regulator of the intestinal villus zonation programme (51). Our data suggests that TGF-β may also play a role in the spatial gene expression programme via ARID3A. The overall increased expression of villus gene signatures across all intestinal epithelial cell types in the mutant animals suggests that ARID3A may have an additional role of spatial differentiation in the villus by fine-tuning the expression levels of zonated genes.

Our data further shows a role of ARID3A in intestinal regeneration. It is believed that progenitor cells in the crypt are highly plastic to allow dedifferentiation into regenerative ISCs upon injury. Deletion of *Arid3a* switches the TA cells to higher differentiation-to-proliferation ratio, which may explain the impaired regenerative capacity by reduced plasticity. This suggests that ARID3A plays a gatekeeping role in the TA compartment to maintain the “just-right” proliferation to differentiation ratio for tissue homeostasis and plasticity. Further investigation of the transcription factor network of ARID3A in the intestinal crypt-villus axis would help understand its unique role in fine-tuning the proliferation-differentiation switch.

## STAR METHODS

### Animals, drug administration and treatments

All animals were maintained with appropriate care according to the United Kingdom Animal Scientific Procedures Act 1986 and the ethics guidelines of the Francis Crick Institute. The full list of transgenic mice used for this study is shown below in Table1.

Animal genotyping was performed by PCR amplification of genomic DNA extracted from ear punch biopsies taken from mice aged three weeks (see primer list on table 2). Biopsies were first digested in 200μl lysis buffer (10mM Tris pH7.5, 100mM NaCl, 10mM EDTA, 0.5% Sarkosyl) at 55oC overnight. 200ng of DNA was amplified using MyTaqTM Red Mix (Bioline, BIO-25043) and 0.4μM forward and reverse primers in a reaction with final volume of 25μl. After the initial denaturation step (95oC for 1min), the thermocycler was configured to 35 cycles of 95oC for 30s, 56-60oC for 30s (depending on the primer) and 72oC for 1min per kilobase of amplification, followed by a final 3min step of extension at 72oC. PCR products were then visualised and size-verified on a 2% w/v agarose/TAE electrophoresis gel with 5ng/ml ethidium bromide. *Villin*Cre-ERT2 animals were genotyped by the Biological Research Facility of the Francis Crick Institute.

To induce conditional deletion, wild-type (WT) and knockout (cKO) animals were injected intraperitoneally with tamoxifen at 1.5mg/10g of mouse weight (from a 20mg/ml stock solution). Mice were culled by schedule 1 procedure (S1K) at the desired timepoint.

For EdU chasing experiments 5-ethynyl-2’-deoxyuridine (EdU) (Life Technologies, E10187) was injected intraperitoneally at 0.3mg/10g of mouse weight (from a 10mg/ml stock solution). Mice were culled by S1K at 2 hr after EdU injection.

For irradiation induced-injury experiments mice were exposed to controlled 12 Gray (12Gy) total body ionising irradiation to induce damage using a Caesium (γ) irradiator. The dosage rate was 0.779Gy/min. Mice were culled by S1K at the desired timepoint.

### Cell culture conditions and maintenance

LS174T cells were maintained in DMEM GlutaMAX (Gibco, 10566-01) supplemented with 5% FBS (Gibco, 10270106) and 100 units/ml penicillin and 100 ug/ml streptomycin (Gibco, 15140122). Cells were incubated in a humidified atmosphere of 5% CO2 at 37°C. For WNT pharmacological inhibition, cells were seeded 24hr before treatment with LGK974 inhibitor (Selleck chemicals, S7143).

### Cell line immunofluorescence

For immunofluorescence (IF) experiments, LS174T cells were grown on sterilised glass coverslips in 24-well plates. Coverslips were not coated with poly-L-lysine before seeding the cells, since LS174T cells are very adherent. Cells were fixed with 4% paraformaldehyde (PFA) for 15min and permeabilised using 0.5% Triton X-100 in PBS for 15min. Cells were blocked with 1% Bovine Serum Albumin (BSA) for 1hr at room temperature before overnight incubation with primary antibodies at 4oC (see table 3 for full list of antibodies). Cells were washed with PBS and incubated with secondary antibodies conjugated to Alexa-Fluor 488 (Invitrogen, A32731) at room temperature for 1hr in the dark. Cells were washed and stained with DAPI and/or Phalloidin-Atto647 (Sigma, 65906) for 30min in the dark. Coverslips were washed and mounted with ProLong Gold Antifade Mountant (ThermoFischer, P36934). Images were acquired as z-stacks using a Leica SPE confocal microscope and processed using Fiji.

### Establishment and maintenance of mouse organoid cultures

Organoids were established from freshly isolated adult small intestine, as previously described (52). In brief, 2cm of jejunal small intestinal tissue was opened longitudinally and villi was scrapped using a glass cover slip. The remaining tissue was incubated in 15mM EDTA and 1.5mM DTT at 4oC for 10min and moved to 15mM EDTA solution at 37oC for an extra 10min. Subsequently, the tissue was shaken vigorously for 30sec to release epithelial cells from basement membrane and the remaining remnant intestinal tissue was removed. Cells were washed once, filtered through a 70μm cell strainer and resuspended in Cultrex BME Type 2 RGF Pathclear (Amsbio, 3533-01002). All freshly isolated organoids were maintained in either Intesticult medium (Stem Cell technologies, #06005) or in-house made basal medium containing EGF (Invitrogen PMG8043), NOGGIN, RSPONDIN and WNT3A (WENR medium), as previously described (28). The Rho kinase inhibitor Y-27632 (Sigma, Y0503) was added to the culture during first week of crypt isolation and single cell dissociation. For WNT, Notch, TGF-β and BMP pathway manipulation, organoids were passaged and allowed to recover for 72 hours before treatment. For WNT signalling inhibition, organoids were treated for 48hr with either 5μM of LGK974 inhibitor or 30μM of LF3 inhibitor (Sigma, SML1752); for Notch signalling inhibition, organoids were treated with 10μM of DAPT inhibitor (Sigma, D5942) for the indicated timepoints; for TGF-β signalling manipulation, organoids were treated with 0.1ng/ml of recombinant TGF-β1 (Sigma, 11412272001) for the indicated timepoints; for BMP signalling manipulation, organoids were treated with 20ng/mL of recombinant BMP4 (Peprotech, 120-05ET) for the indicated timepoints.

NOGGIN and RSPONDIN conditioned media were generated by HEK293T cells. WNT3A conditioned medium was generated from L cells. All images were acquired using an EVOS FL Cell Imaging System (Life technologies) and image brightness was adjusted using Adobe Photoshop (exactly same parameters were applied to all samples of the same experiment).

### Mouse organoids assays

Organoids were established as described in the previous section. For organoid formation assay, crypts were counted using a brightfield microscope and 200 crypts were seeded in 20μl of Cultrex BME Type 2 RGF Pathclear in individual wells of a 48-well plate and cultured in WENR medium for 5 days until counted. Three technical replicates were performed per animal.

For RSPONDIN withdrawal assay, organoids were passaged and seeded in 3 10μl droplet per well of a 24-well plate. Organoids were allowed to re-establish in normal ENR medium (5% RSPONDIN) for 48hr. Subsequently, organoid medium was replaced and have organoids were cultured in ENR medium containing either 5% or 1% of RSPONDIN.

To detect disaccharide levels in organoids supernatants, organoids were washed twice with PBS and incubated with a 56mM solution of sucrose during 1h. Supernatants were collected and frozen until the assay was performed. To detect glucose content, Amplex® Red Glucose/Glucose Oxidase Assay Kit (Invitrogen, A22189) was used. Samples were diluted when necessary and incubated with the reaction buffer containing Amplex Red®, horseradish peroxidase and glucose oxidase. Fluorescence was measured in a Tecan microplate reader with an excitation wavelength of 540nm and fluorescence emission detection at 590nm. Glucose concentration was assessed using a glucose standard curve from 0 to 200µM.

### Crypt-villus fractionation

4cm of jejunal small intestinal tissue was opened longitudinally and villi was scrapped using a glass cover slip. Villi and the remaining intestinal tissue were transferred into two separate tubes and washed once. Both parts of the small intestine were incubated in 15mM EDTA and 1.5mM DTT at 4oC for 10min and moved to 15mM EDTA solution at 37oC for an extra 10min (as described in section 2.3.1) for isolation of epithelial cells of villi and crypt fragments. Pelleted cells were re-suspended in RLT buffer and stored at -800C before proceeding to RNA extraction.

### Fluorescent-activated Cell Sorting (FACS) of GFP-positive cells from *Lgr5-EGFP-ires-CreERT2 mice*

Crypts were harvested from the proximal jejunum (∼10cm) as described in section “Establishment and maintenance of mouse organoid cultures”. Crypts were then dissociated by incubating with Collagenase/Dispase (Roche, 11097113001) for 20min at 37oC, followed by 20min incubation with TrypLE (Gibco, 12604013) for 20min at 37oC. TrypLE was stopped by adding Advanced DMEM (Gibco, 12491015) containing 10% fetal bovine serum (FBS) (Gibco, 10270106) and dissociated cells were passed through a 20μm strainer. Cells were stained with DAPI and resuspended in PBS-0.5% BSA-2mM EDTA. Cells were separated and re-collected in Advanced DMEM plus 10% FBS based on GFP intensity. Cell sorting was performed on a BD FACSAria™ II System.

### RNA isolation from cell lines, organoids and tissue

RNA was extracted according to the manufacturer’s instructions (Qiagen RNeasy, 74106). Harvested cell lines, organoids or intestinal crypts were resuspended in RLT buffer (provided with the kit) supplemented with 40mM Dithiothreitol (DTT) to inhibit RNases activity. Before RNA extraction samples were undergone one freeze/thaw cycle to increase the yield of extracted RNA.

### cDNA synthesis and qRT-PCR

500-1000ng of RNA were reverse transcribed using the cDNA Reverse Transcription Kit (Applied Biosystems, #4368813), according to manufacturer’s instructions. RT-qPCR was performed in 384-well plates, in experimental triplicates, in a 12μl reaction mixture containing 6μl of 2x PowerUp™ SYBR® Green Master Mix (Applied Biosystems, A25742), 10μM of each primer and 25-50ng of cDNA. The reaction mixture without cDNA template was used as a negative control for each reaction plate. After 40 cycles of amplification, samples were normalised to housekeeping genes *Ppib* (for mouse samples) or *β-ACTIN* (for human samples), where data was expressed as mean ± s.e.m.

### Immunohistochemistry

For analysis of small intestine by immunohistochemistry (IHC), tissues were fixed in 10% formalin and embedded in paraffin. Sections were deparaffinized with xylene and rehydrated in a graded series of ethanol. Antigen-retrieval was performed for 20 min at high temperature in o.01M Citrate (pH 6) or Tris-EDTA buffer (10mM Tris base, 1mM EDTA, pH 9). Slides were then blocked using the appropriate blocking buffer (10% Normal goat serum or 10% Normal donkey serum in 1% BSA) and incubated overnight with the appropriate antibody at 4°C (See table 3 for full list of antibodies). Finally, slides were incubated with the secondary antibody for 1h and washed three times with PBS. For colorimetric staining, with diaminobenzidine (DAB) slides were incubated with peroxidase substrate, dehydrated, counterstained with Hematoxylin solution according to Mayer (Sigma, 51275) and mounted. Slides were scanned using an Olympus VS120 slide scanner and images were processed using QuPath (53).

For immunofluorescence, slides were incubated with Alexa-Fluor 488 or Alexa-Fluor 568 antibody for 1h, washed three times with PBS, incubated with 4’,6’-diamidino-2-phenylindole (DAPI) for 15 min to visualize nuclear DNA and mounted with ProLong Gold Antifade Mountant. Images were acquired as z-stacks using a Leica SPE or a Leica SP8 confocal microscope and processed using Fiji. For whole slide imaging, slides were scanned using an Olympus VS120 slide scanner and images were processed using QuPath (53).

When indicated, sections were stained for Hematoxylin & Eosin (H&E), alkaline phosphatase and Alcian Blue-Periodic Acid Schiff (AB-PAS) staining. Edu was detected according to the manufacturer’s protocol (Thermo Fisher Scientific, Click-iT Plus EdU Alexa Fluor 555 imaging kit C 10638) to evaluate proliferating cell number. Edu+ cells were quantified in at least 10 crypts per mouse.

### *In situ* hybridisation

Single-molecule *in situ* hybridization was performed on mouse intestine according to manufacturer’s instructions (ACD; RNAscope 2.5HD Assay RED (REF 322350) or RNAscope 2.5HD (REF 322436)). The probes used were against *Arid3a* (REF 525721), *Arid3b* (REF 525731), *Atoh1* (REF 408791), *Lgr5* (REF 312171) and *Olfm4* (REF 311831). Briefly, guts were fixed in formalin overnight, paraffin-embedded and cut into 5-micron thick slices. Target retrieval was performed for 15 minutes, followed by RNAScope Protease Plus incubation for 24 minutes on the FFPE Sample Preparation and subsequent amplification steps. For brightfield analysis, slides were counterstained using 50% Hematoxylin solution according to Mayer and for immunofluorescence slides were incubated with DAPI for 10min for DNA visualisation. Images acquired with an Olympus VS120 slide scanner and images were processed using QuPath (53).

For combined RNAscope and immunofluorescence of *Arid3a* with MUC2, CHGA or LYZ, samples were first stained for *Arid3a* using the red channel of duplex RNAscope kit, followed by antibody immunostaining as described above.

**Table 1.** List of antibodies.

| ANTIBODY | APPLICATION | DILUTION | MANUFACTURER |
| --- | --- | --- | --- |
| ARID3A | Cell line IFA | 1:100 | Abcam (169787) |
| ARID3A | IHC | 1:700 | Proteintech (14068-1-AP) |
| MUC2 | IHC | 1:200 | Santa Cruz (sc-15334) |
| CHGA | IHC | 1:200 | Abcam (ab15160) |
| LYZ | IHC | 1:2000 | Dako (A0099) |
| ASPA | IHC | 1:100 | Proteintech (13244-1-AP) |
| APOA4 | IHC | 1:500 | R&D (AF8125) |
| CLEAVED CASPASE -3 | IHC | 1:200 | Cell signalling (9664) |
| KI67 | IHC | 1:200 | Abcam (ab15580) |
| CD235-APC | FACS | 1:100 | Biolegend (118214) |
| CD31-PE | FACS | 1:100 | Biolegend (102407) |
| CD45-FITC | FACS | 1:100 | Biolegend (157214) |

### Bulk RNA-seq sample preparation

Crypts or villi were isolated from 10cm of mouse jejunal small intestinal tissue as described in “Crypt-villus fractionation” section and RNA was isolated as described in “RNA isolation” section. RNA integrity (RIN) was examined using Bioanalyzer 2100 RNA 6000 Nano kit from Agilent and RIN cut-off was set to 7. For crypt samples, libraries were prepared using KAPA mRNA HyperPrep kit (KK8580) according to manufacturer’s instructions. For most villus samples, RIN number was lower than 7 and libraries were prepared with KAPA RNA HyperPrep with RiboErase (KK8561) according to manufacturer’s instructions.

### Bulk RNA-seq data analysis

Fastq files were processed using the nf-core/RNASeq pipeline (10.5281/zenodo.4323183) version 3.0 using the corresponding versions of STAR RSEM to quantify the reads against release 95 of Ensembl GRCm38. These raw counts were then imported into R (R Core Team (2020). R: A language and environment for statistical computing. R Foundation for Statistical Computing, Vienna, Austria. URL https://www.R-project.org/) version 4.03 /Bioconductor version 3.12 (54). We then used to DESeq2 (55)version 1.30.1 to account for the different size factors between the samples, and used a generalised linear (negative binomial) model with main effects of arid3a (or arid3b) status and time (as a categorial variable) to find genes that were statistically significantly associated with arid3a (or arid3b) status (Wald test) with a false discovery rate of <0.05.

Gene set enrichment analysis (GSEA) was performed using the GSEA desktop software (version 4.1.0) using the following parameters: GSEA Preranked > no collapse of gene symbols > classic enrichment statistic > Chip platform “Mouse_Gene_Symbol_Remapping_Human_Orthologs_MSigDB.v7.5.chip”. GSEA custom lists were obtained from the indicated publications. For Metacore analysis, the online software was used (https://portal.genego.com/). Gene lists of upregulated and downregulated genes were created by using FDR<0.05 and FC>1.5 cut-offs. One click analysis included Pathway Maps and GO Processes.

### ATAC-seq sample preparation

Isolated mouse crypts were dissociated to single cells as described in section “FACS of GFP-positive cells from *Lgr5-EGFP-ires-CreERT2* mice*”* and single cell numbers and viability were assessed using Trypan blue dye and Neubauer chamber. 25.000 cells per sample were transferred to a fresh tube, pelleted and incubated in RSB buffer (10mM Tris-Cl pH 7.4, 10mM NaCl and 3mM MgCl2) supplemented with 0.1% v/v NP-40 (Sigma, 11332473001), 0.1% v/v Tween-20 (Sigma, 11332465001) and 0.01% Digitonin (Promega, G9441) in order to isolate intact nuclei. Isolated nuclei were subsequently treated with Tn5 transposase (Illumina, 20034197) for 30min at 37oC with agitation for DNA tagmentation. DNA was immediately purified using Qiagen Mini Elute kit (Qiagen, 28004). Purified DNA was subsequently used for library preparation using NEBNext® High-Fidelity 2X PCR Master Mix (NEB, M0541S) using primers with Nextera dual indexes in a 20μl final reaction volume. PCR amplification included 5min incubation at 72oC, followed by 30sec of DNA denaturation at 98 oC and 12 cycles of the following: 98 oC for 10sec, 63 oC for 30sec and 72 oC for 1min. PCR products were cleaned-up using Ampure XP beads (Beckman Coulter, A63881) according to manufacturer’s instructions. The quality of the final DNA library was confirmed on the Agilent Tapestation before the samples were submitted to sequencing.

### ATAC-seq data analysis

The nf-core/atacseq pipeline (version 1.2.1) (Ewels et al., 2020) written in the Nextflow domain specific language (version 19.10.0) (Di Tommaso et al.,2017) was used to perform the primary analysis of the fastq samples in conjunction with Singularity (version 2.6.0) (Kurtzer et al., 2017). The command used was “nextflow run nf-core/atacseq -profile crick --input /Path_to_desing/design.csv –fasta Mus_musculus.GRCm38.dna_sm.toplevel.fa - -gtf Mus_musculus.GRCm38.95.gtf --gene_bed Mus_musculus.GRCm38.95.bed --macs_gsize 2.6e9 --blacklist mm10.blacklist.bed --narrow_peak -r 1.2.1 -resume”.

To summarise, the pipeline performs adapter trimming (Trim Galore!- https://www.bioinformatics.babraham.ac.uk/projects/trim_galore/), reads alignment (BWA) and filtering (SAMtools) (56), (BEDTools) (57); BamTools (58); pysam - https://github.com/pysam-developers/pysam; picard-tools - http://broadinstitute.github.io/picard)), normalised coverage track generation ((BEDTools) (57); bedGraphToBigWig (59)), peak calling (MACS) (60) and annotation relative to gene features (HOMER) (61), consensus peak set creation (BEDTools), differential binding analysis ((featureCounts) (62) R Core Team, DESeq2 (55)) and extensive QC and version reporting ((MultiQC (63), FastQC (64), deepTools (65), ataqv (66)). All data was processed relative to the mouse Ensembl GRCm38 release 95. A set of consensus peaks was created by selecting peaks that appear in at least one sample. Counts per peak per sample was then imported on DESeq2 within R environment for differential expression analysis. Pairwise comparisons between genotypes in each condition, and between conditions per genotype were carried out and differential accessible peaks were selected with an FDR < 0.05.

For footprinting analysis TOBIAS (v 0.12.10) (35) was used by running the following pipeline (https://github.com/luslab/briscoe-nf-tobias). The pipeline runs TOBIAS’ ATACorrect, ScoreBigwig, BINDetect and generates PlotAggregate metaplots on merged replicate bam files. TOBIAS was run on set of consensus peaks used for the differential analysis (see above). As described before (35), all TFs with -log10(p-value) above the 95% quantile or differential binding scores smaller/larger than the 5% and 95% quantile are coloured. Selected TFs are also shown with labels.

For ChromVar analysis, the previously published R package was used: (http://www.github.com/GreenleafLab/chromVAR), to analyse sparse chromatin-accessibility data by estimating gain or loss of accessibility within peaks sharing the same motif or annotation while controlling for technical biases. Identified TFs were sorted based on their variability score.

For GO analysis, the 5000 peaks with the highest variability between WT and *Arid3a* cKO samples were chosen, and peaks were annotated with the closest gene prior to the analysis.

### scRNA-seq sample preparation

Isolated mouse crypts were dissociated to single cells as described in section “FACS of GFP-positive cells from *Lgr5-EGFP-ires-CreERT2* mice*”* and single cell numbers and viability were assessed using Trypan blue dye and Neubauer chamber. 3106 cells per sample were stained for CD45-FITC (Biolegend, 103108), Epcam-APC (Biolegend, 118214) and CD31-PE (Biolegend, 102507) at 1/100 dilution during 30min at RT in the dark. DAPI was added at a 1/1000 dilution for the last 5min. After antibody incubation and washing, samples were re-suspended in FACS buffer (PBS-0.5% BSA-2mM EDTA). Single cell stainings and unstained controls were used for gating. Cells were sorted for DAPI-negative to select for live cells, Epcam-positive to acquire epithelial cells, CD45-negative to exclude lymphocytes and CD31-negative to exclude endothelial cells. After sorting, cells were pelleted and re-suspended in 1000cells/μl final concentration and cell viability was assessed as a quality check parameter. Only samples with viabilities higher tan 85% were processed for library preparation, which was done using 10X_3primer_mRNA as per manufacturer’s instructions. Samples were sequenced using Novaseq platform. A minimum of 5000k reads per cell was reached and paired-end sequencing was performed.

### scRNA-seq data analysis

Raw sequencing data was processed using the CellRanger pipeline (version 5.0; 10X Genomics) (67). Count tables were loaded into R (version 4.0.3) and processed using the Seurat R-package for single-cell analysis (version 4.0.5) (68). All cell with fewer than 1000 distinct genes observer per cell and cells with more than 15% of unique molecular identifiers stemming from mitochondrial genes were removed as low-quality cells prior to the downstream analysis. Integration of individual samples was done using the canonical correlation analysis method from the Seurat package with 3000 integration features. Principal component analysis was performed on the 2000 most variable genes in the dataset and the first 30 principal components were selected as input for UMAP dimensionality reduction and clustering. Clustering was performed with the default method of the Seurat R-package, with the resolution parameter set to 0.3. Cluster marker genes were inspected and corresponding cell types assigned manually. Single-cell differential gene expression analyses were performed using the R-package glmGamPoi (version 1.2.0) (69). Trajectory analyses were conducted using the monocle3 R-package (version 0.2.2) (70) with the ISC cluster 0 set as starting point. A sub-clustering of cells in the ISC and TA clusters was performed. For the sub-clustering analysis, the cluster resolution parameter was set to 0.3 and clusters were assigned manually. Cell cycle phases were estimated using the CellCycleScoring function of the Seurat R-package.

## Acknowledgments

We thank Joan Yuan (Lund University) and Stephen Malin (Karolinska Institute) for providing the Arid3a floxed mice. We also thank the Francis Crick Institute’s Biological Research Facilities, Experimental Histopathology, Advanced Sequencing facilities, Bioinformatics and Biostatistics for technical supports. This work was supported by the Francis Crick Institute, which receives its core funding from Cancer Research UK (CC2141), the UK Medical Research Council (CC2141), and the Wellcome Trust (CC2141).

## Data and code availability

The accession numbers for the raw and processed data used for this study was deposited on GEO and is publicly available (Bulk RNA-seq GEO: GSE242983, scRNA-seq GEO: GSE243155 and ATAC-seq GEO: GSE212560).

## FIGURE LEGENDS

**Supplementary figure 1.**
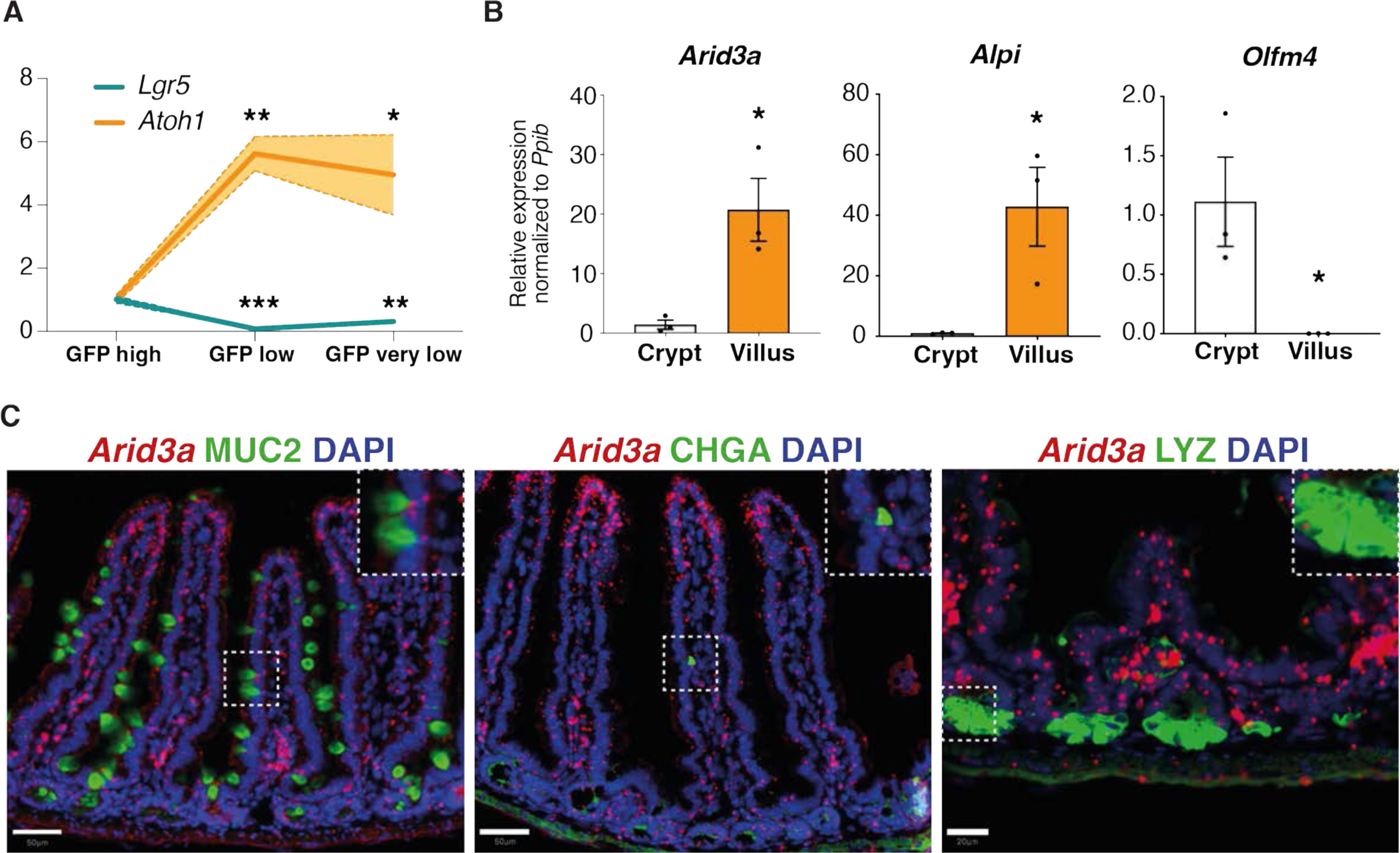
*Arid3a* has higher expression in the villus compared to the crypt. (A) qRT-PCR analysis of three FACS sorted populations representing ISCs (GFP high) and progenitor cells (GFP low and GFP very low). Three biologically independent animals (N=3). Data represent mean ± s.e.m. (light colour area). (B) qRT-PCR analysis of crypt and villus fractions. Three biologically independent animals (N=3) *P<0.05, **P<0.01, ***P<0.001, two-sided t-test. (C) Combined *Arid3a* RNAscope scope staining with immunofluorescent protein staining of MUC2, CHGA and LYZ. Each staining was performed at three different animals (N=3). Scale bar, 50μm for *Arid3a*/MUC2 and *Arid3a*/CHGA and 20μm for *Arid3a*/LYZ.

**Supplementary figure 2.**
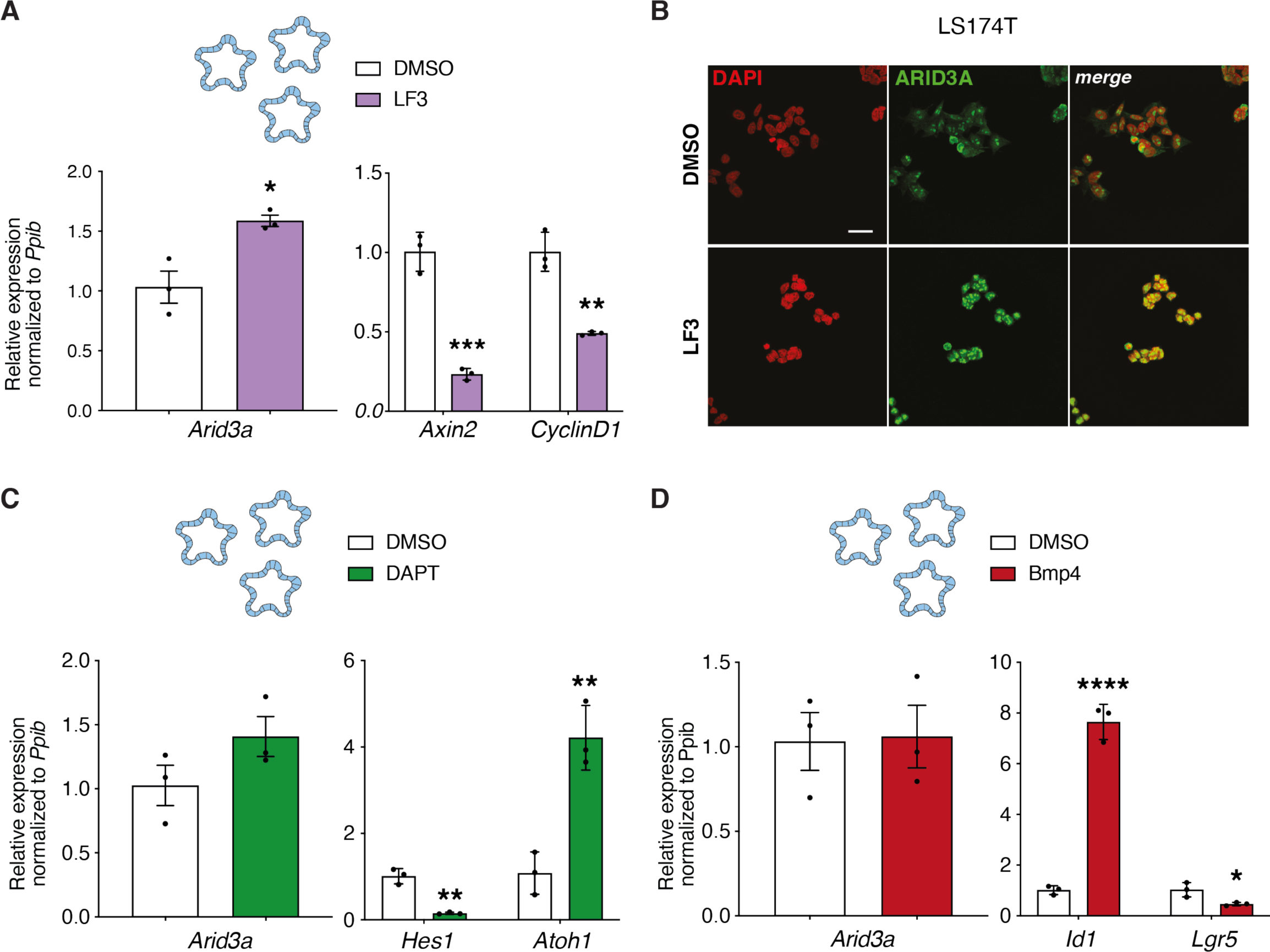
BMP and NOTCH signalling do not affect *Arid3a* expression. (A) qRT-PCR analysis of WT organoids treated with LF3 inhibitor for 48h. Organoids were established from three biologically independent animals per group (N=3). (B) Immunofluorescence staining for ARID3A upon LF3 treatment of LS174T cells, scale bar 100μm. Three independent experiments were performed (n=3). (C) qRT-PCR analysis of WT organoids treated with DAPT inhibitor for 48h. Organoids were established from three biologically independent animals per group (N=3). (D) qRT-PCR analysis of WT organoids treated with recombinant BMP4 for 4h. Organoids were established from three biologically independent animals per group (N=3). Data represent mean ± s.e.m. *P<0.05, **P<0.01, ***P<0.001, two-sided t-test.

**Supplementary figure 3.**
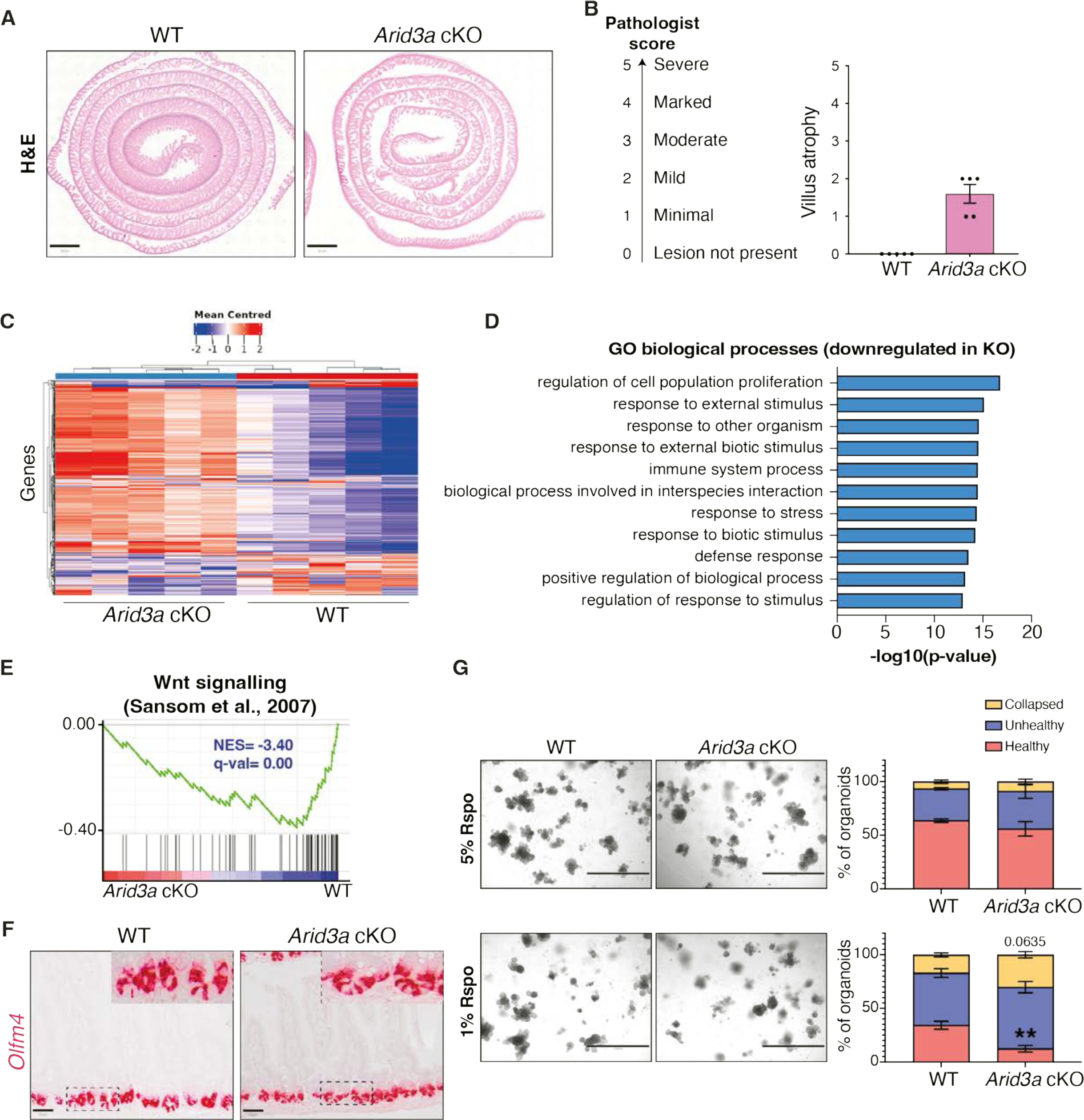
*Arid3a* cKO animals have reduced proliferative capacity. (A) H&E staining of WT and *Arid3a* cKO mice. Representative images of N=5 animals per experimental group. Scale bar, 100μm. (B) Results of the pathologist’s report of WT and *Arid3a* cKO small intestinal tissue. N=5 animals per experimental group. (C) Heatmap of blinded transcriptome-wide gene expression. The colours in the heatmap are a gene’s expression in a sample, relative to its average expression. (D) Top 10 downregulated GO biological processes based on RNA-seq analysis. FDR<0.05 and fold change> 1.5 cut-offs were applied. (E) GSEA of previously published WNT signalling gene list. (F) RNAscope staining of *Olfm4* in WT and *Arid3a* cKO mice. Representative picture of N=3 per group. Scale bar, 50μm. (G) Representative images and quantification of WT and *Arid3a* cKO organoids cultured in 5% or 1% Rspo CM. N=8 WT and N=4 Arid3a cKO organoid lines were used. Stacked data represent mean ± s.e.m. *P<0.05, **P<0.01, ***P<0.001, 2-way ANOVA.

**Supplementary figure 4.**
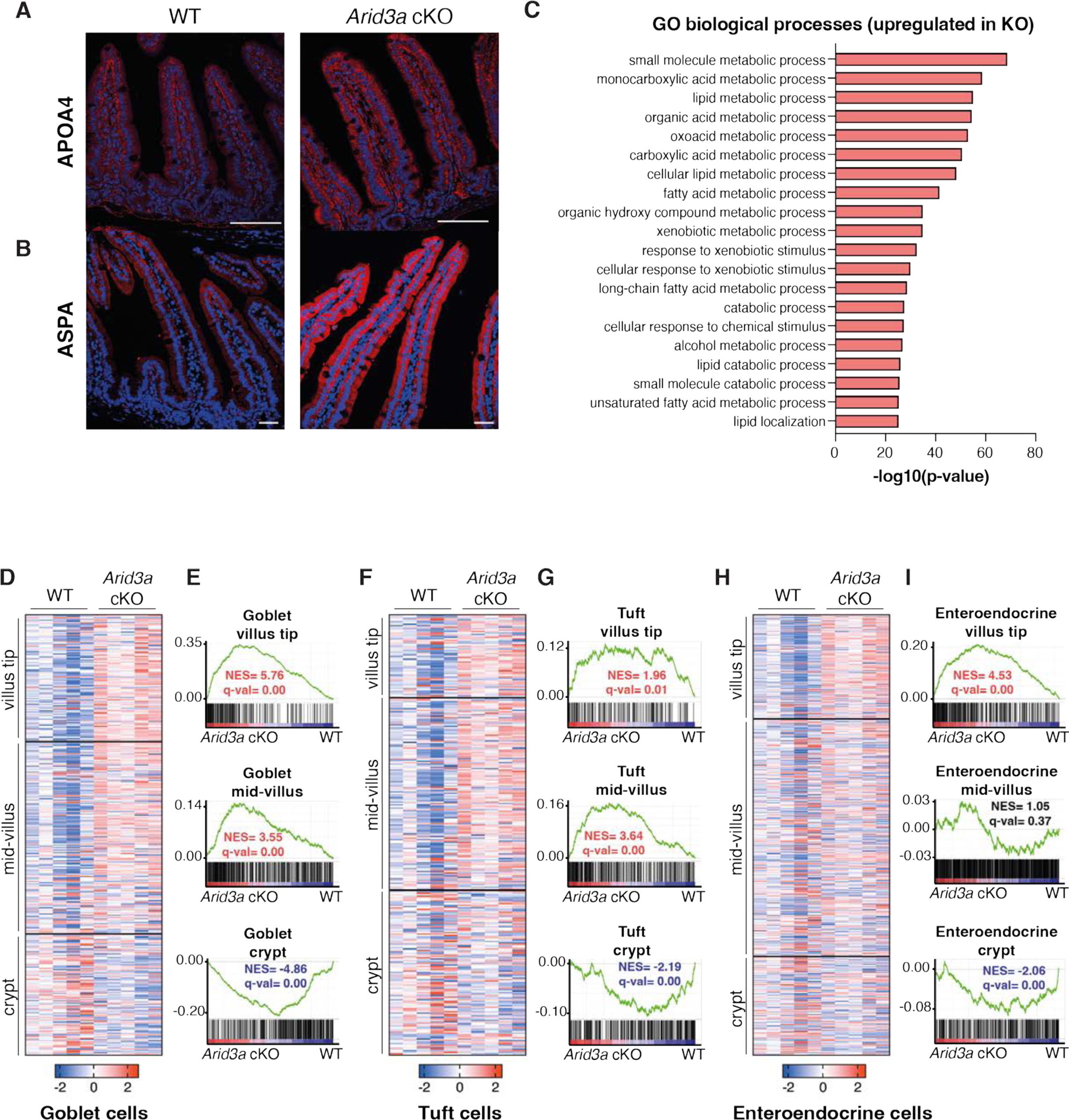
Loss of *Arid3a* leads to disruption of enterocyte zonation. (A) Immunofluorescence staining of APOA4. Representative images from N=5 per experimental group. Scale bar, 100μm. (B) Immunofluorescence staining of ASPA. Representative images from N=3 per experimental group. Scale bar, 100μm. (C) Top 20 upregulated GO biological processes based on RNA-seq analysis. FDR<0.05 and fold change> 1.5 cut-offs were applied. (D) Heatmap of RNA-seq data of goblet cell zonated and differentially expressed genes. Z-scores are shown. FDR cut-off <0.05. (E) GSEA of all zonated goblet cell genes. (F) Heatmap of RNA-seq data of tuft cell zonated and differentially expressed genes. Z-scores are shown. FDR cut-off <0.05. (G) GSEA of all zonated tuft cell genes. (H) Heatmap of RNA-seq data of enteroendocrine cell zonated and differentially expressed genes. Z-scores are shown. FDR cut-off <0.05. (I) GSEA of all zonated enteroendocrine cell genes. For (G), (I) and (K) genes are shown based on their centre of mass with crypt genes at the bottom of the heatmap and villus tip genes at the top (based on Manco et al., 2021).

**Supplementary figure 5.**
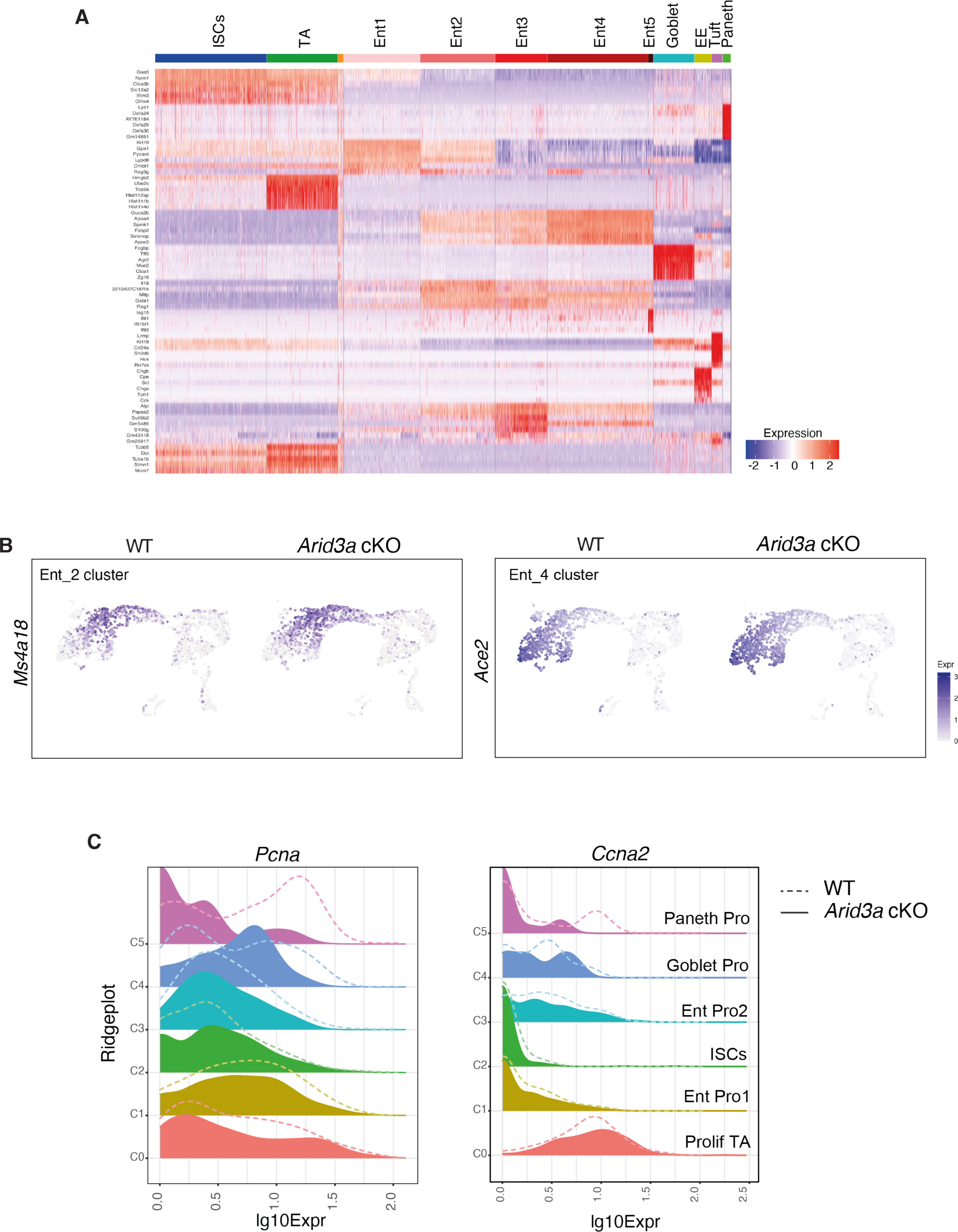
Single-cell RNA-sequencing reveals changes in enterocyte differentiation trajectory and cell cycle phase in TA cells. (A) Top Cluster Markers Heatmap showing the most distinct marker genes in each cluster. Cluster annotation was based on identification of previously published genes within this list. Cluster annotated in orange colour included a very small number of cells with no specific enrichment for specific genes. This cluster was excluded from any further downstream analysis. (B) Feature plot of *Ms4a18* (enterocyte marker cluster 2) and *Ace2* (enterocyte marker cluster 5) expression on UMAP. (C) Ridgeplots showing log10 expression of *Pcna* and *Ccna2* genes in WT and *Arid3a* cKO for each crypt cell sub-population.

**Supplementary figure 6.**
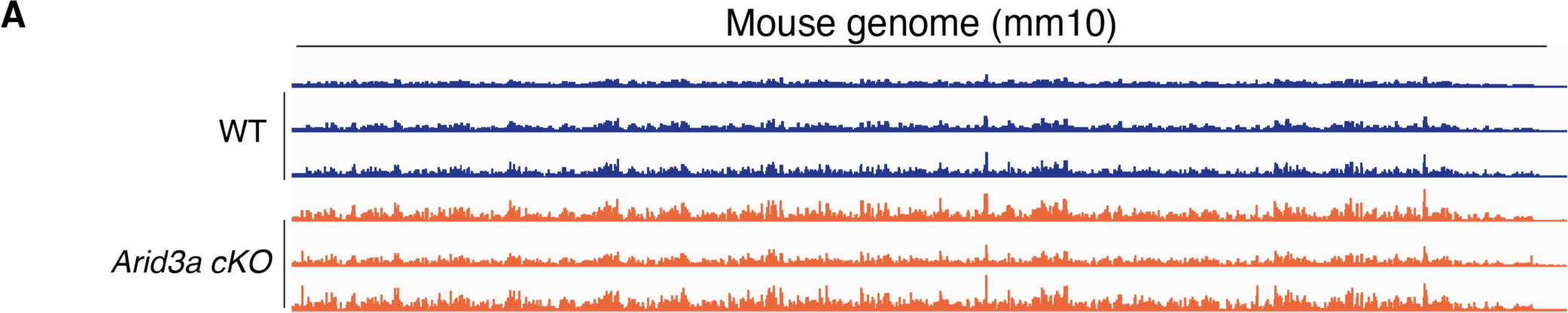
Deletion of Arid3a results in genome-wide open chromatin. (A) Genome-wide comparison of open ATAC-seq chromatin peaks between WT and *Arid3a* cKO animals (tracks extracted from IGV).

**Supplementary figure 7.**
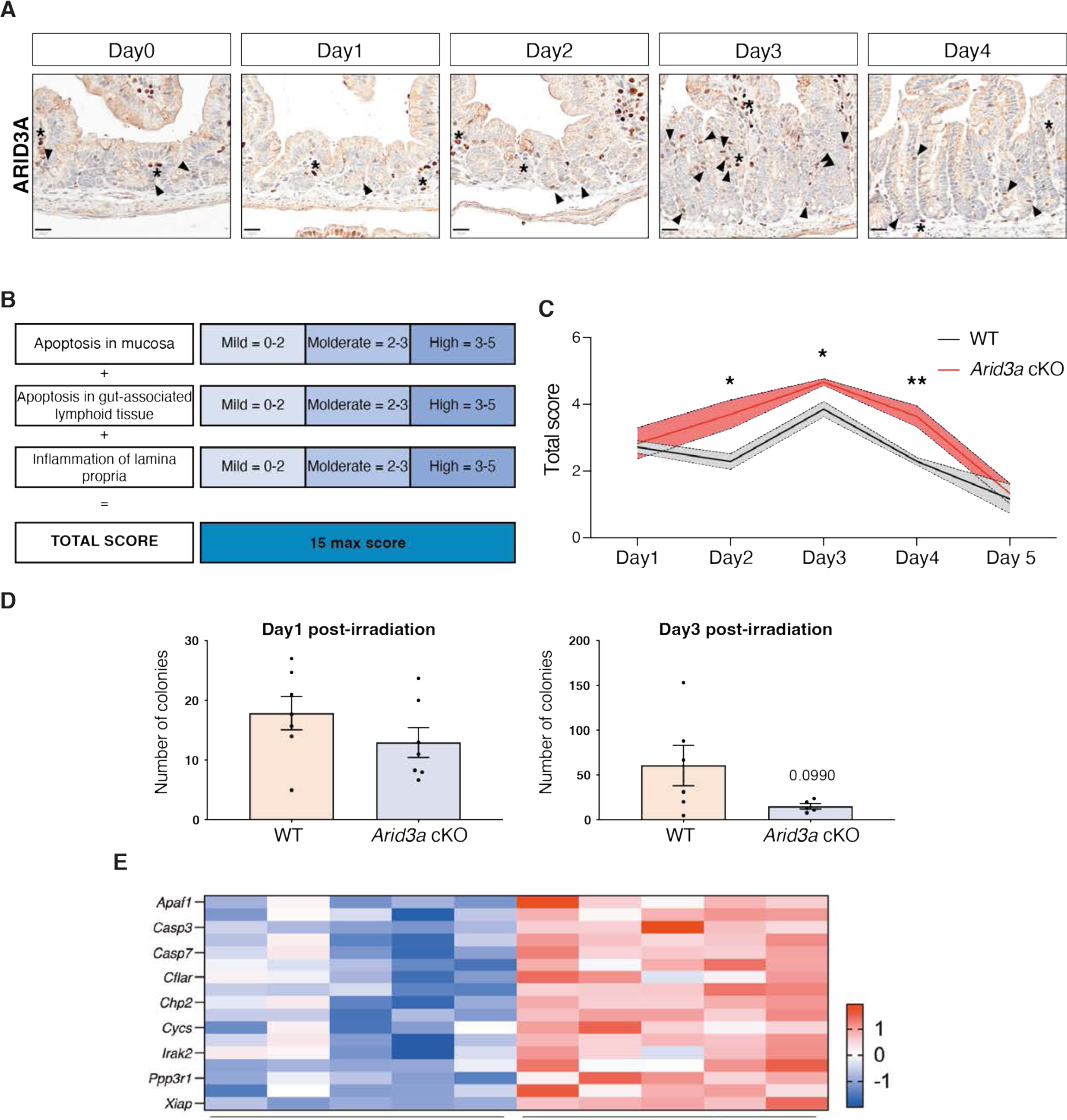
Loss of Arid3a leads to reduced proliferation and extended tissue damage after irradiation. (A) ARID3A immunostaining of WT mice during the first 4 days after administration of 12Gy irradiation. Representative images of N=5 WT animals per timepoint. Scale bar, 20μm. Black arrowheads indicate epithelial expression of ARID3A; asterisks indicate stromal expression of ARID3A. (B) Description of the scoring system used to assess damage of the small intestine at days. (C) Quantification of tissue damage. Data represent mean ± s.e.m. (light colour area) *P<0.05, **P<0.01, ***P<0.001, two-sided t-test. (D) Organoid formation assay was performed at days 1 and 3 post irradiation. Data represent mean ± s.e.m. *P<0.05, **P<0.01, ***P<0.001, two-sided t-test. (E) Heatmap of RNA-seq data of representative cell death-associated markers. Z-scores are shown.

## SUPPLEMENTAL INFORMATION

**Supplemental information 1 |** Differential gene expression analysis of Arid3a cKO crypts versus WT

**Supplemental information 2 |** Cluster specific gene signatures based on scRNA-seq analysis

